# Identifying Explainable and Generalizable Features for MEG Decoding

**DOI:** 10.1101/2025.04.30.651376

**Authors:** Nima Maleki, Hamid Karimi-Rouzbahani

## Abstract

Sensory neural coding, the brain’s process of transforming inputs into informative patterns of neural activity, generates complex and multiplexed neural codes which are hard to interpret. Although decoding methods have facilitated the interpretation of these codes, the specific features of neural activity that generalize across individuals, along with their precise timing, largely remain elusive. To address this gap, we investigated the potential of interpretable time-series features in magnetoencephalography (MEG) for decoding visual stimulus attributes (spatial frequency and orientation) and their generalizability across individuals. We extracted a comprehensive set of highly informative features from 18 subjects engaged in a visual task and performed a time-resolved decoding analysis. Our findings revealed that particular features, especially those capturing rapid changes in neural activity within the first 200 milliseconds of stimulus presentation, yielded high decoding accuracy. Notably, these early, transient features exhibited robust cross-subject generalizability, suggesting a shared neural coding mechanism during the initial processing of visual inputs. Furthermore, these features outperformed previously successful electroencephalography (EEG) wavelet features when applied to MEG data. While within-subject decoding demonstrated sustained above-chance performance, cross-subject generalization diminished after the initial 200 milliseconds, indicating more individualized processing at later stages. Our results underscore the importance of systematic, data-driven evaluation of neural signals for elucidating neural codes and for developing more transparent and generalizable Brain-Computer Interface (BCI) systems that capitalize on these reliable neural signatures.

## Introduction

The process of information coding by neurons is often referred to as neural coding, where different types of information (e.g., sensory, motor and cognitive) are converted into patterns of spiking neuronal activity. Different aspects of these spiking activities such as their precise timing and frequency encode different types of information (Borst & Theunissen, 1996; Panzeri et al., 2010). For example, hippocampal place cells’ average firing rate conveys information about animal’s spatial position, creating spatial maps of the environment (Poucet et al., 2004). Understanding these encoding mechanisms reveals how neural coding supports complex brain processes in health and disease and real-world applications such as BCIs which bridge the gap between neural activity and artificial prostheses (Barbieri et al., 2004).

Neural coding is supported by complex interactions between neuronal assemblies (Harris, 2005) and synaptic plasticity (Feldman, 2012). Synaptic connections form patterns of neural activity as permitted sets, shaped by excitatory and inhibitory interactions. These mechanisms support learning and memory, as in the hippocampal place neurons, which can reconstruct patterns from limited inputs (Curto et al., 2013). Neural assemblies further encode information by forming selectively activated groups of neurons that propagate signals, supporting functions like memory and language. For example, retinal ganglion cells illustrate how sensory systems refine input, compressing visual information for efficient perception (Kartsaki et al., 2024; Papadimitriou & Friederici, 2022). The combination of these encoding mechanisms produces remarkably rich, complex and often multiplexed neural codes (Panzeri et al., 2010; Karimi-Rouzbahani, 2024), which have been partially decoded using a variety of methods in the literature (Auge et al., 2021; Hancock & Khoshgoftaar, 2020; Paninski et al., 2007; Vargas-Hakim et al., 2021). The development of such decoding pipelines continues to date (van Gerven et al., 2019; Koide-Majima et al., 2024).

Neural decoding aims at translating neural activity into interpretable information. It has been extensively explored in various domains, including motor control, sensory perception, and cognitive processing. For instance, studies have shown that neural decoding models can effectively intercept and decode movement intentions by analyzing brain signals from the primary motor cortex in humans (Tam et al., 2019). Similarly, sensory domain studies have revealed that neural responses in the visual cortex can be decoded to reveal information about object orientation and color, suggesting robust encoding schemes in sensory processing areas (Miyawaki et al., 2008; Graf et al., 2011). Neural decoding methods encompass a broad spectrum, ranging from classical statistical and mathematical approaches like Markov and Bayesian models (Ahmadian et al., 2011; R Barbieri et al., 2004; Barbieri et al., 2005; Endres et al., 2009; Kemere et al., 2003; Oram et al., 1998; van Gerven et al., 2019) and Kalman Filters (Bishop et al., 2008; Malik et al., 2010; Wu et al., 2009), to advanced machine learning techniques, particularly deep learning methods (Glaser et al., 2020; Santos-Mayo et al., 2025; Zhang et al., 2022) each with successful applications to different datasets. These methods have demonstrated significant improvements in decoding complex neural signals and have not only enhanced our basic understanding of brain function but have also enabled applications like BCIs (Wolpaw et al., 2002). Moreover, the development of novel feature extraction techniques, where one focuses on informative aspects of recorded neural activity (ignoring non-informative aspects), has helped to capture essential temporal and spectral characteristics of neural data, thereby further improved the robustness of decoding algorithms (Rybakken et al., 2019; Karimi-Rouzbahani et al., 2017; Karimi-Rouzbahani et al., 2021). These advancements have paved the way for neural decoding applications in various neuroimaging modalities in humans.

Neural decoding in humans has dominantly been applied to non-invasive neuroimaging modalities such as functional Magnetic Resonance Imaging (fMRI), electroencephalography (EEG) and magnetoencephalography (MEG) to attribute specific patterns of brain activity to specific task conditions including sensory (Haxby et al., 2001; Kay et al., 2008; Naselaris et al., 2009), motor (Hatsopoulos & Donoghue, 2009; Schalk et al., 2004; Wolpaw et al., 2002), and cognitive tasks (Blankertz et al., 2011; Grootswagers et al., 2017; Moerel et al., 2021). While fMRI has superior spatial resolution compared to EEG and MEG, the latter modalities have higher temporal resolution and can provide millisecond-scale insights into the temporal dynamics of information encoding in the brain (Baillet, 2017; Hämäläinen et al., 1993; Karimi-Rouzbahani & McGonigal, 2024; King & Dehaene, 2014; Siegel et al., 2012). EEG, as a more common and accessible neural recording modality, has attracted the application of many decoding pipelines including time-resolved decoding (Carlson et al., 2019). However, MEG, while also providing great temporal resolution, has received less attention in neural decoding. This is an important gap as MEG captures neural activities which are produced by neuronal sources orthogonal to the ones captured by EEG (Gross, 2019), providing complementary insights into the dynamics of information processing in the brain. Moreover, with the growing use of mobile MEG systems such as optically pumped magnetometers (OPMs; Hill et al., 2024) in both basic (Schofield et al., 2024) and applied neuroscience domains (Wittevrongel et al., 2021), more rigorous MEG decoding algorithms are needed to harness its full potential.

While MEG decoding has been used to answer many neuroscientific questions (Baillet, 2017; Gross, 2019; Hari & Salmelin, 2012; Kutas & Dale, 2013), it remains unknown how generalizable the neural codes produced in MEG are from one brain to another. In other words, it is unclear if two human brains produce similar neural codes for undertaking the same cognitive process. For example, we do not know if one subject encodes an image in a similar way to another. Moreover, the precise timing of cross-subject generalizable neural codes remains under-investigated. The latest MEG decoding studies have generally performed the decoding analysis for each individual subject and used a single feature of neural activity (e.g., signal mean) in decoding (Hebart et al., 2018; Mohsenzadeh et al., 2019; Rajaei et al., 2019; Karimi-Rouzbahani et al., 2024). This leaves us wonder if there are specific patterns of neural activity which generalize across subjects while others do not. We have tried to evaluate the role of several features of neural activity for neural decoding in non-invasive (Karimi-Rouzbahani, 2024), and invasive (Karimi-Rouzbahani & McGonigal, 2024; Karimi-Rouzbahani & McGonigal, 2025) EEG modalities. Specifically, we have evaluated the information content of a large set of features extracted in EEG and found that multiscale patterns of activity, as quantified through multiscale Wavelet coefficients, provided the maximum decoding performance (Karimi-Rouzbahani et al., 2021). However, such multi-featural approach is lacking in the MEG domain, which has its own distinct spatiotemporal characteristics to EEG leading to an under-explored wealth of MEG data to be explored. More importantly, cross-subject generalizability of informative neural codes in MEG has not yet been systematically investigated. We know from years of research that neural activity patterns vary across individuals, which can hinder the performance of individualized decoding methods when generalizing across subjects (Miller et al., 2002; Wei et al., 2021). Therefore, to develop more effective decoding methods, we need to identify the features of neural activity which generalize across subjects.

While cross-subject generalization has been a focus of recent studies using transfer learning and domain adaptation techniques (Csaky et al., 2023; Wang et al., 2024), little is known about the generalizable features of MEG neural codes and their timings as they have used black-box deep learning approaches. To fill this gap, the current study has developed a data-driven, explainable decoding approach to identify the features of neural activity which generalize across distinct portions of data within an individual subject. Specifically, we extracted 22 distinct features of neural activity, which have been systematically selected from a large set of 7,700 features (Lubba et al., 2019). In our dataset of 18 MEG subjects, we evaluated which features generalized across subjects and if the *within-subject* and *cross-subject* sets of generalizable features correlated.

## Methods

### MEG Dataset

We used an MEG dataset previously collected using a visually evoked retro-cue working memory task, as detailed in (Quentin et al., 2019)^1^. The original dataset included data from 22 subjects, where each subject participated in two separate computer-based experimental sessions. Each session comprised recordings from 272 MEG channels, and the original signals were collected at 1200Hz sampling frequency and down-sampled to 120Hz, which was the version we used in the present study.

The dataset initially included MEG recordings from 22 subjects, each with two sessions divided into eight runs. However, preprocessing revealed issues with four subjects, where some runs had small data sizes—approximately 10% of the typical run size—causing errors in analysis. To maintain consistency, these four subjects were excluded, leaving a final dataset of 18 subjects, each with two complete sessions.

### Experimental Task and Stimuli

We provide a brief explanation of the task (Quentin et al., 2019). The task was performed by subjects while MEG was collected, with visual stimuli back-projected onto a translucent screen positioned in front of them. The experimental task presented visual gratings varied in spatial frequencies (1, 1.5, 2.25, 3.375, or 5.06 cycles/°), and orientations (−72°, −36°, 0°, 36°, and 73°, with 0° representing vertical) making 25 possible combinations of spatial frequencies and orientations (Figure 1A). Each trial started with a fixation dot on the center of the screen, which subjects were instructed to focus on throughout the trial (Figure 1B). After a jittered interval of 400 ms (±50 ms), two semi-circled visual gratings were presented simultaneously for 100 ms, one in each half of the visual field. The two gratings were never identical. While each trial also involved a cue and probe after the second fixation period, in the present study we only analyzed the visual stimulus presentation part of the trial as shown in Figure 1B.

**Figure 1.**
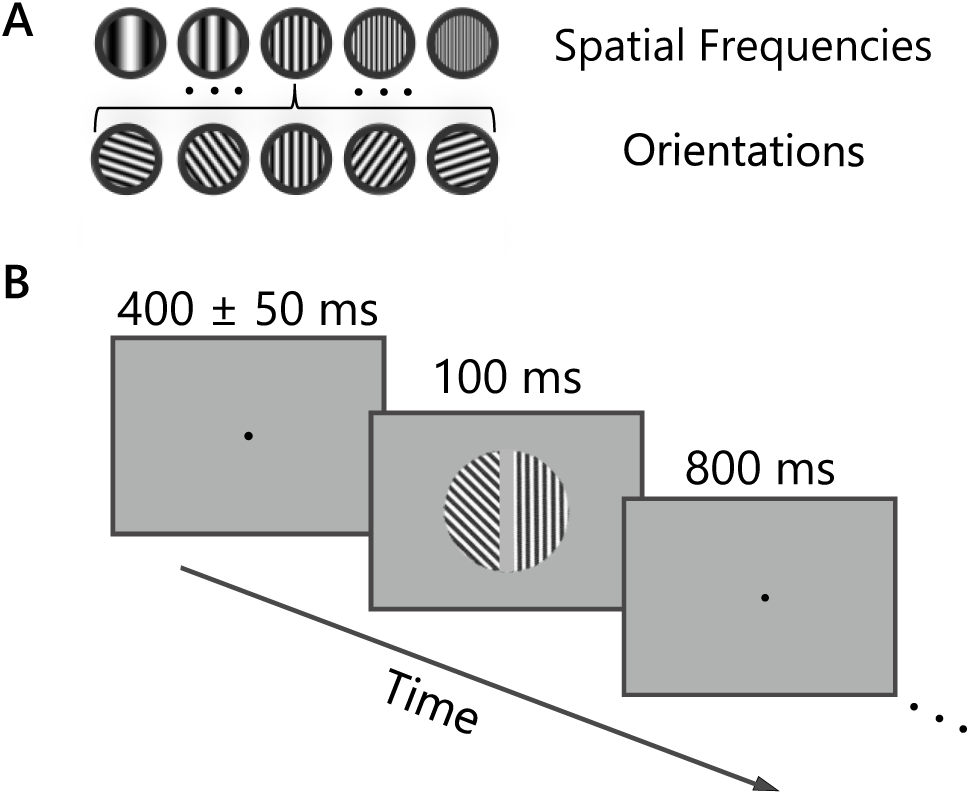
Stimuli and task. **A** shows the 5 different spatial frequencies and 5 orientations of the grating stimuli which were presented to subjects. **B** shows (stimulus presentation part of) the trial where subjects were presented with a fixation dot followed by two grating stimuli and a follow-up fixation dot with the indicated timings. Figure reconstructed from Quentin et al., 2019.

### Signal Preprocessing

The MEG data had been band-pass filtered in the range 0.05–25 Hz and then decimated by a factor of 10, resulting in a final sampling frequency of 120 Hz. We segmented the signals into epochs from −0.2s before to 0.9s after the stimulus onset and baselined them between −0.2s and 0s relative to the stimulus onset. The data from the two MEG sessions for each subject were concatenated for analyses.

### Feature Extraction

In our analysis, we initially extracted a comprehensive set of 25 distinct features from neural time series to investigate both informative and generalizable patterns of neural activity across subjects. Each of these features was used separately in decoding to allow for assessing their information content for decoding and their ability to generalize across different subjects. This methodological approach allowed direct comparison across features without being affected by their interactions and provided valuable insights into both *within-* and *cross-subject* decoding of neural data.

Of these 25 features, 24 were derived from the updated version of the Catch22 (Lubba et al., 2019), (Canonical Time-Series Characteristics) feature set, which has been carefully selected to efficiently quantify informative aspects of time-series data. Originally, Catch22 extracted 22 canonical time-series characteristics from the much larger hctsa (Highly Comparative Time-Series Analysis) repository (Fulcher & Jones, 2017), which extracts approximately 7,700 time-series features. These 22 features were carefully selected to represent a wide range of statistical, spectral, and temporal properties that are informative in the context of time-series analysis (Lubba et al., 2019). In the updated version of Catch22, two additional features—“mean” and “standard deviation”—were included, further enhancing the feature set’s capacity to summarize key aspects of the data. This makes the Catch22 feature set a powerful tool for summarizing complex time-series data, such as the neural signals recorded in our study.

In addition to the Catch22 features, we also included a wavelet-based feature in our analysis. Wavelet transforms are widely used in signal processing for their ability to capture both time and frequency domain information simultaneously, making them particularly well-suited for analyzing non-stationary signals like those found in MEG recordings. Wavelet features were added as they provided the highest performance among the features in EEG decoding previously (Taghizadeh-Sarabi et al., 2015; Karimi-Rouzbahani et al., 2021) and we wanted to compare the current Catch22 features with the wavelet feature in MEG neural decoding.

However, three features from Catch22 features, were excluded from further analysis:

The two features of *SC FluctAnal2 rsrangefit_50_1_logi_prop_r1* and *SC FluctAnal2 dfa_50_1_ 2_logi_prop_r1* required a window size of a minimum of 50 samples for meaningful analysis, so were excluded from analyses. Our whole trial epoch had 133 samples, and using 50 of them would lead to the loss of the temporal information we were interested in. Additionally, the feature *PD_PeriodicityWang_th0_01* performed near the chance level in analyses and was excluded. This feature has been designed to measure periodicity based on Wang’s method (Wang et al., 2007) and the low periodicity in our dataset seem to have led to its chance-level performance. Therefore, it was excluded from our analyses. After excluding these three features, we proceeded with a final set of 22 features.

Through the application of these 22 distinct features, we aimed to gain a deeper understanding of which features were the most informative for decoding neural activity and which were more generalizable across different subjects, contributing to the overall goal of improving neural decoding performance. Features were extracted using 10-sample partially overlapping sliding windows over the course of the trial.

Below, we briefly describe each of the features used in this study, which were obtained from Canonical Time-Series Characteristics^2^ and the wavelet feature we used previously in EEG (Karimi-Rouzbahani et al., 2021). For feature extraction, we utilized the PyCatch22^3^ and PyWT (PyWavelets for wavelet transform).

For ease of interpretation, we have categorized our features into eight groups based on their working mechanisms and conceptual frameworks, which are detailed below.

### Distribution Shape (DB Shape)

These features assess distribution shapes of time-series values, focusing on how values are distributed over time and identifying frequent values. They help understand the data’s form and central tendencies, revealing distinctive patterns. The Catch22 toolkit implements two such features using the DN_HistogramMode function as detailed below.

#### DN_HistogramMode_5 (mode_5)

This feature calculates the mode of the z-scored time series using a histogram with 5 linearly spaced bins. The mode is the value that appears most frequently in the data set. By z-scoring the time series, the data is normalized to have a mean of zero and a standard deviation of one, making the mode calculation more robust to variations in scale and location. For example, the mode in the 5-binned histogram reveals the most frequent brain activity level in a given time window of MEG activity.

#### DN_HistogramMode_10 (mode_10)

Similar to DN_HistogramMode_5, this feature calculates the mode of the z-scored time series but uses a histogram with 10 linearly spaced bins. This provides a finer resolution of the data distribution, allowing for a more detailed analysis of the most frequent values in the time series, but can be noisier with shorter time series. After z-scoring MEG data, the mode in a 10-binned histogram reveals the most frequently occurring brain activity level, revealing finer details compared to DN_HistogramMode_5.

### Extreme Event Timings (Extreme)

These features are designed to measure the timing of extreme events within a time series, specifically looking at how these events occur in relation to the beginning and end of the series. By doing so, they help identify when significant outliers happen over time. The Catch22 toolkit includes two features that utilize the DN_OutlierInclude function, providing valuable insights into the temporal distribution and impact of these extreme events in time-series data.

#### DN_OutlierInclude_p_001_mdrmd (outlier_timin_pos)

This feature measures the median range of time series with a 0.001 probability threshold, assessing outliers’ impact on distribution and variability. It standardizes the time series, analyzes over-threshold events/samples using z-scores, and computes their median index rescaled from −1 to 1. This feature reflects the impact of outliers by measuring the median range as more outliers are included. It identifies whether these events are concentrated near the beginning, evenly distributed, or towards the end of the time series.

#### DN_OutlierInclude_n_001_mdrmd (outlier_timin_neg)

Similar to DN_OutlierInclude_p_001_mdrmd, this feature quantifies the median of the time series as more outliers are included, but with a negative probability threshold of 0.001. This feature assesses the influence of outliers on the time series. This feature measures how negative outliers affect MEG data. It shows whether these events cluster near the start, spread evenly, or gather towards the end of the time series.

### Linear Autocorrelations (Linear Auto)

The linear autocorrelation structure features quantify the relationships between values in a time series based on their linear dependencies. These features are derived from the autocorrelation function or the power spectrum, helping to identify and measure how past values in the series influence future values. This quantification aids in understanding the persistence and predictability of patterns within the data. This category includes four features as follows.

#### CO_f1ecac (acf_timescale)

This feature measures the first lag of the autocorrelation function of the time series. Autocorrelation is the correlation of a signal with a delayed copy of itself, and the first lag indicates how much the current value of the series is related to its immediate past value.

#### CO_FirstMin_ac (acf_first_min)

This feature identifies the first minimum of the autocorrelation function of the time series. The first minimum represents the first point where the autocorrelation drops to a local minimum, providing insights into the periodicity and structure of the time series. This feature pinpoints the periodic patterns and the overall structure of neural activity in the MEG data.

#### SP_Summaries_welch_rect_area_5_1 (low_freq_power)

This feature computes the area under the PSD curve using Welch’s method. It calculates relative power in the lowest 20% of frequencies, assigning higher values to significant low-frequency power and lower values to high-frequency power. The PSD is estimated within a linear space.

#### SP_Summaries_welch_rect_centroid (centroid_freq)

Similar to the previous feature, this feature is also derived from the power spectrum, estimated using Welch’s method with a rectangular window. However, in this case, it identifies the frequency, f, at which the power in frequencies below and above f is equal, referred to as the ‘centroid’. This feature calculates the MEG data’s PSD, indicating the frequency where power distribution is balanced. High values suggest concentration in higher frequencies.

### Nonlinear Autocorrelations (Nonlinear Auto)

The nonlinear autocorrelation features are designed to capture the properties of a time series that involve nonlinear dependencies. Unlike linear autocorrelation, which measures direct, proportional relationships, these features help identify and quantify more complex, nonlinear relationships between values over time. This category includes three features as follows.

#### CO_trev_1_num (trev)

This feature quantifies the time-reversibility of the time series by comparing the number of increases and decreases in the series. A time series is time-reversible if its statistical properties are invariant under time reversal. This feature helps in identifying asymmetries in the time series.

#### CO_HistogramAMI_even_2_5 (ami2)

This feature calculates the average mutual information (AMI) of the time series using a histogram with 5 bins and a lag of 2. AMI measures the amount of information shared between a time series and its lagged version, indicating the level of dependency (memory) between them. We selected a lag of 2 samples in the current work, quantifying the short-delayed dependency in neural signals.

#### IN_AutoMutualInfoStats_40_gaussian_fmmi (ami_timescale)

This feature measures the first minimum of the auto mutual information function of the time series, using a Gaussian kernel with a lag of 40 as the default in the toolbox. Auto mutual information quantifies the amount of shared information between a time series and its lagged version, helping to identify temporal dependencies and patterns.

### Symbolic

The symbolic features are derived from converting real-valued time-series data into discrete symbols. This approach helps in analyzing the data by summarizing it into a sequence of symbols, making it easier to identify patterns, trends, and anomalies within the time series. This symbolic representation can capture important aspects of the data’s behavior in a more manageable form, providing valuable insights into its structure and dynamics. This category includes four features as follows.

#### SB_MotifThree_quantile_hh (entropy_pairs)

This feature detects recurring patterns of length three in the time series using a high-high quantile threshold. It converts the series into three symbols, analyses two-letter sequence probabilities, and outputs entropy, with low values indicating higher predictability and high values indicating lower predictability. This feature spots how often a specific pattern of length three appears in the MEG data using high-high quantile thresholds. It translates the series into symbols to track and analyze recurring patterns, revealing their predictability.

#### SB_TransitionMatrix_3ac_sumdiagcov (transition_variance)

This feature computes the sum of the diagonal covariance of a 3×3 transition matrix from the time series, indicating the stability of state transitions. It converts the series into a 3-letter alphabet and computes τ-step transition probabilities as a 3×3 matrix, returning the sum of column-wise variances. Low values suggest uniform transitions, while high values suggest ordered transitions. higher values of this feature suggest more predictable patterns of signal activity from one window of time to the next.

#### SB_BinaryStats_mean_longstretch1 (stretch_high)

This feature identifies the longest sequence of consecutive values in a time series that exceeds the average. It converts the series into a binary sequence (values above the avearge as 1, others as 0) and returns the longest run of 1s. This feature finds the longest sequence of consecutive data points highlighting extended periods of heightened activity.

#### SB_BinaryStats_diff_longstretch0 (stretch_decreasing)

This feature calculates the length of the longest stretch of consecutive decreases in the time series. It provides insights into the persistence of downward trends and the overall stability of the series. It converts the series into a binary sequence (1 for increases, 0 for decreases) and returns the longest run of 0s. This feature identifies the longest series of consecutive decreases in MEG data, reflecting periods of decline and assesses stability.

### Incremental Differences (Incremental)

The incremental differences features capture the properties of the time series by looking at the differences between consecutive points. By examining these 1-point incremental changes, these features provide insights into the rate of change and the variability within the series, helping to identify trends, patterns, and potential anomalies in the data. This category has two features as follows.

#### MD_hrv_classic_pnn40 (high_fluctuation)

This feature assesses the proportion of differences between consecutive data points that exceed 4% of the standard deviation of the time series, distinguishing between variability patterns. Low values indicate minimal fluctuations within 0.04 times the standard deviation, while high values indicate significant jumps. It highlights the contrast between steady periods and abrupt changes.

#### FC_LocalSimple_mean1_tauresrat (whiten_timescale)

This feature calculates the ratio of the mean of the time series to the mean of residuals after fitting a local model with a time delay, providing information on predictability and smoothness. It measures prediction errors using the mean of the previous 3 values in a z-scored series and standard deviation of 1-step forecast residuals. This feature evaluates sample predictability by comparing the mean of the time series to the mean of residuals after a simple local model fit.

### Simple Forecasting (Forecasting)

The simple forecasting features focus on generating forecasts based on the mean of the previous three points in the time series. This method provides a straightforward way to predict future values by averaging the recent past values, offering insights into short-term trends and helping to smooth out minor fluctuations in the data. This category has only one feature as follows.

#### FC_LocalSimple_mean3_stderr (forecast_error)

This feature computes the standard error of the mean after fitting a local model with a time delay, providing information about variability and reliability. It measures prediction errors using the mean of the previous 3 values in a z-scored series and standard deviation of 1-step forecast residuals. This feature assesses the variability and reliability of MEG data, after fitting a local model with a time delay.

### Moments and Embedding Dist

Moment features are statistical measures that provide essential insights into the distribution of time-series data. These features help summarize and describe the fundamental characteristics of the data, allowing for the identification of patterns, trends, and anomalies. This category has two features as follows.

#### DN_Mean (mean)

This feature calculates the mean value of the time series. The mean is a measure of central tendency, representing the average value of the data points in the series. This feature calculates the average value of the MEG data points, providing a central tendency of the neural activity over the recorded period.

#### DN_Spread_Std (std)

This feature calculates the standard deviation of the time series. The standard deviation is a measure of dispersion, indicating how spread out the data points are around the mean. It shows how much the brain activity deviates from the mean value over the recording period.

#### CO_Embed2_Dist_tau_d_expfit_meandiff (embedding_dist)

This feature calculates the mean difference between distances in an embedded space with a time delay and a dimension (d), offering insights into temporal structure and complexity. It maps the time series onto a 2D time-delay embedding space, computes distances, and analyzes their probability distribution, outputting the mean absolute error of an exponential fit. Low values indicate a good fit to the exponential distribution. In the MEG data’s temporal structure, low values suggest a strong exponential fit, indicating the data’s complexity and patterning.

#### Wavelet

The wavelet feature excels in time-series analysis, especially for non-stationary signals like MEG data. Unlike Fourier analysis, it represents signals in both time and frequency domains, capturing transient events and localized frequency changes. The wavelet transform decomposes the signal into scaled and shifted versions of a wavelet function, enabling multi-resolution analysis and detecting patterns at different scales. This feature is crucial in identifying time-varying neural dynamics, often missed by single-domain features. Widely used in neuroscience (Taghizadeh-Sarabi et al., 2015; Karimi-Rouzbahani et al., 2021), the wavelet feature captures both short-lived and long-lasting neural patterns.

### Time-Resolved Decoding

We used a sliding window time-resolved decoding analysis to quantify the discriminability of grating stimulus attributes. Specifically, we used the features extracted from the sliding windows (i.e., with a width of 10 samples/80ms with 5 samples/40ms overlap), which allowed fine-grained decoding of stimulus attributes. Our data captured neural activity evoked by grating visual stimuli each having two attributes: spatial frequency (i.e., how dense the lines were within a given area) and orientation (i.e., angle of lines relative to the horizontal line). Each attribute had five categorical conditions (five spatial frequencies and five orientations). We made binary classifications where we systematically decoded all possible pairwise combinations of stimuli with a given attribute, resulting in 10 unique binary classifications per stimulus attribute. For example, we decoded gratings of 0° from 36° and 0° from 72° and so on, irrespective of spatial frequency and decoded gratings with a spatial frequency of 1 from 1.5 cycles/° and 1 from 2.25 cycles/° and so on, irrespective of orientations. We averaged the classification scores across these 10 combinations to obtain robust and comprehensive decoding metrics. Figure 2 shows our time-resolved neural decoding analysis pipeline.

**Figure 2.**
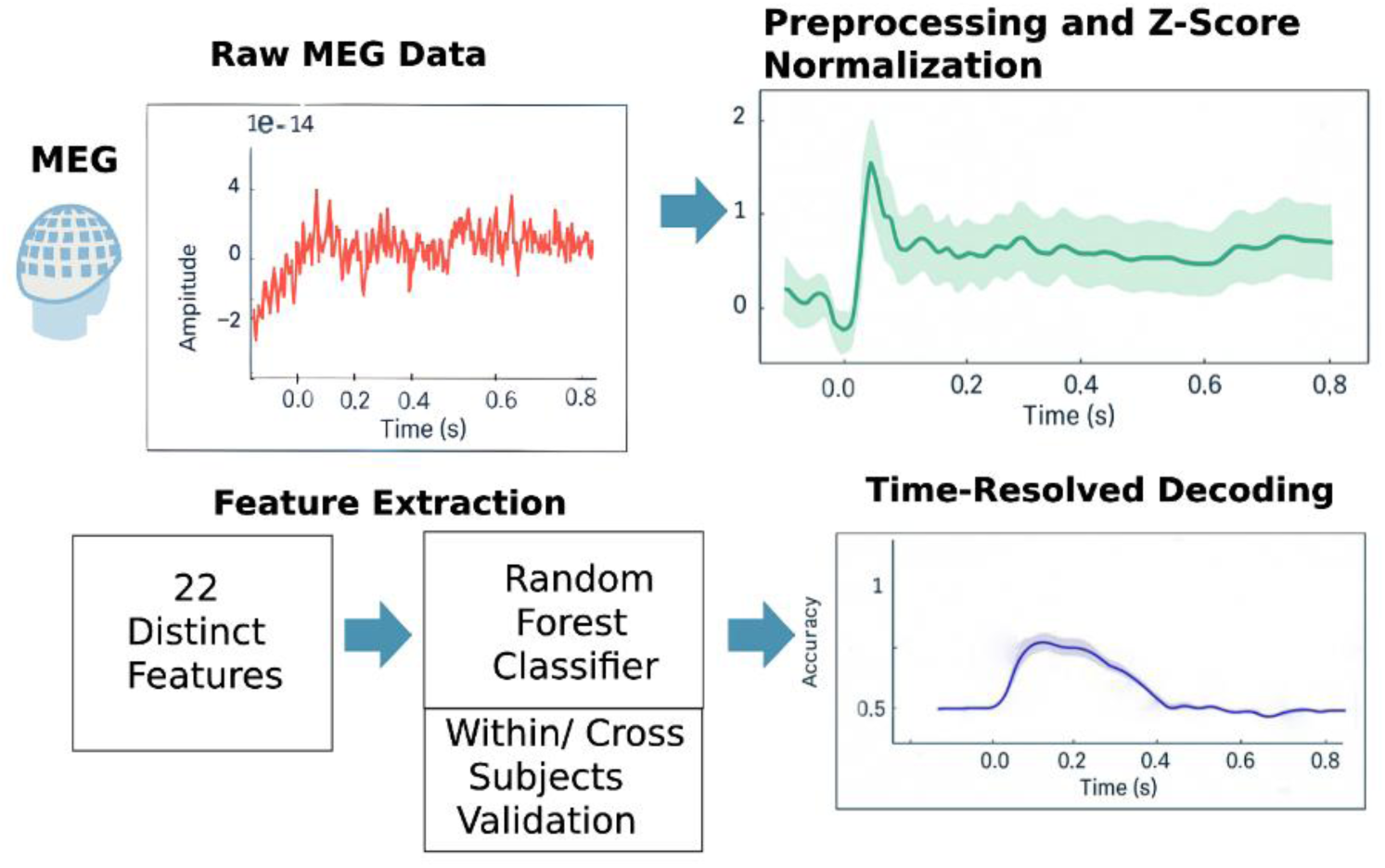
Time-resolved MEG decoding pipeline. The pipeline starts with preprocessing the MEG signals (including filtering and normalization), feature extraction followed by the application of classification algorithms to decode neural activity over time.

### Random Forest Classifier

We used a Random Forest classifier for decoding as in our previous work (Karimi-Rouzbahani & McGonigal, 2024). It is a robust ensemble learning method widely used for its versatility and resistance to overfitting (Fernández-Delgado et al., 2014). It constructs multiple decision trees using bootstrapped data subsets, with random feature selection at each split to enhance generalization. The final prediction is obtained by combining the outputs of all decision trees, taking the mode of the classes for classification or the mean value for regression. Known for handling large, noisy datasets effectively, Random Forest has been extensively applied in neuroscience to decode neural data and classify cognitive states (Breiman, 2001; Chen et al., 2014). To implement our classifier, we used the Scikit-Learn library in Python.

### Within-Subject Analysis

*Within-subject* analysis assessed feature generalizability within the same subject. By testing how well the features perform across different portions of a subject’s dataset, we can determine the consistency and reliability of the features for that individual.

In the *within-subject* analysis, we used a 12-fold cross-validation scheme to evaluate the decoding performance in out-of-sample data. This means that each subject’s dataset was divided into 12 equal parts or “folds”. For each iteration, one fold of data was left out for testing, while the remaining 11 folds were used for training the model. This process was repeated 12 times, with each fold serving as the test set exactly once.

While checking the generalizability of patterns within a subject, the *within-subject* method does not test the ability of the features to generalize across different subjects, as the training and testing were confined to a single subject’s dataset.

### Cross-Subject Analysis

*Cross-subject* analysis assessed the generalizability of features across subjects. Specifically, it measured how well the features learned from one group of subjects could be informative about an unseen subject. This is critical for understanding the generalizability of features in applications where models need to work on new subjects not included in the training dataset. This method ensures the model captures patterns which are generalizable to unseen subjects while reducing overfitting to subject-specific patterns (Chen et al., 2014).

The *cross-subject* analysis employed a Leave-One-Subject-Out (LOSO) cross-validation approach. Our dataset consisted of data from 18 subjects. In each iteration, the data from one subject was left out from the training set and used as the test set, while the data from the remaining 17 subjects was used to train the model. This process was repeated 18 times until each subject served as the test set once.

### Correlation Analysis

We used *Pearson* correlation to examine the statistical relationships between *within-subject* and *cross-subject* analyses. Pearson correlation measures the strength and direction of a linear relationship between two variables. This helped us evaluate the consistency and generalizability of our features across individuals (Cohen et al., 2009). For the sake of presentations, we also fitted lines to the data.

### Statistical Analysis

#### Bayes Factor

We utilized Bayes factor analysis as a statistical tool to evaluate the evidence for one hypothesis over another, particularly when comparing decoding results against a baseline level of decoding. The Bayes factor is a ratio of the likelihoods of two competing hypotheses, typically the null hypothesis (no effect) and the alternative hypothesis (presence of an effect). A Bayes factor greater than 1 indicates evidence in favor of the alternative hypothesis, while a value less than 1 supports the null hypothesis. Importantly, the Bayes factor offers a more nuanced measure than traditional p-values, as it quantifies the degree of evidence rather than simply rejecting or accepting a hypothesis. This makes it particularly useful in neuroscience and decoding studies, where the goal is often to assess the relative strength of observed effects. To interpret the Bayes factor results, we chose strict thresholds and considered evidence above 6 as evidence in favor of the alternative hypothesis, and values below 1/6 indicating strong evidence for the null (Karimi-Rouzbahani, 2024).

## Results

We conducted a time-resolved decoding analysis to investigate how well specific features of neural activity could discriminate/decode stimulus attributes (i.e., different spatial frequencies or orientations of grating stimuli) and whether these features were generalizable across individuals. Within the manuscript, we present only the decoding results for spatial frequency and the orientation decoding results are presented in the supplementary materials.

### Within-Subject Analysis

First, we performed neural decoding using one feature of neural activity at a time to discriminate the stimuli at different spatial frequencies. Figure 3 presents the decoding results for each feature and feature category and compares them to the decoding results obtained from the *wavelet* feature which outperformed other features in earlier EEG decoding studies (Taghizadeh-Sarabi et al., 2015; Karimi-Rouzbahani et al., 2021).

**Figure 3.**
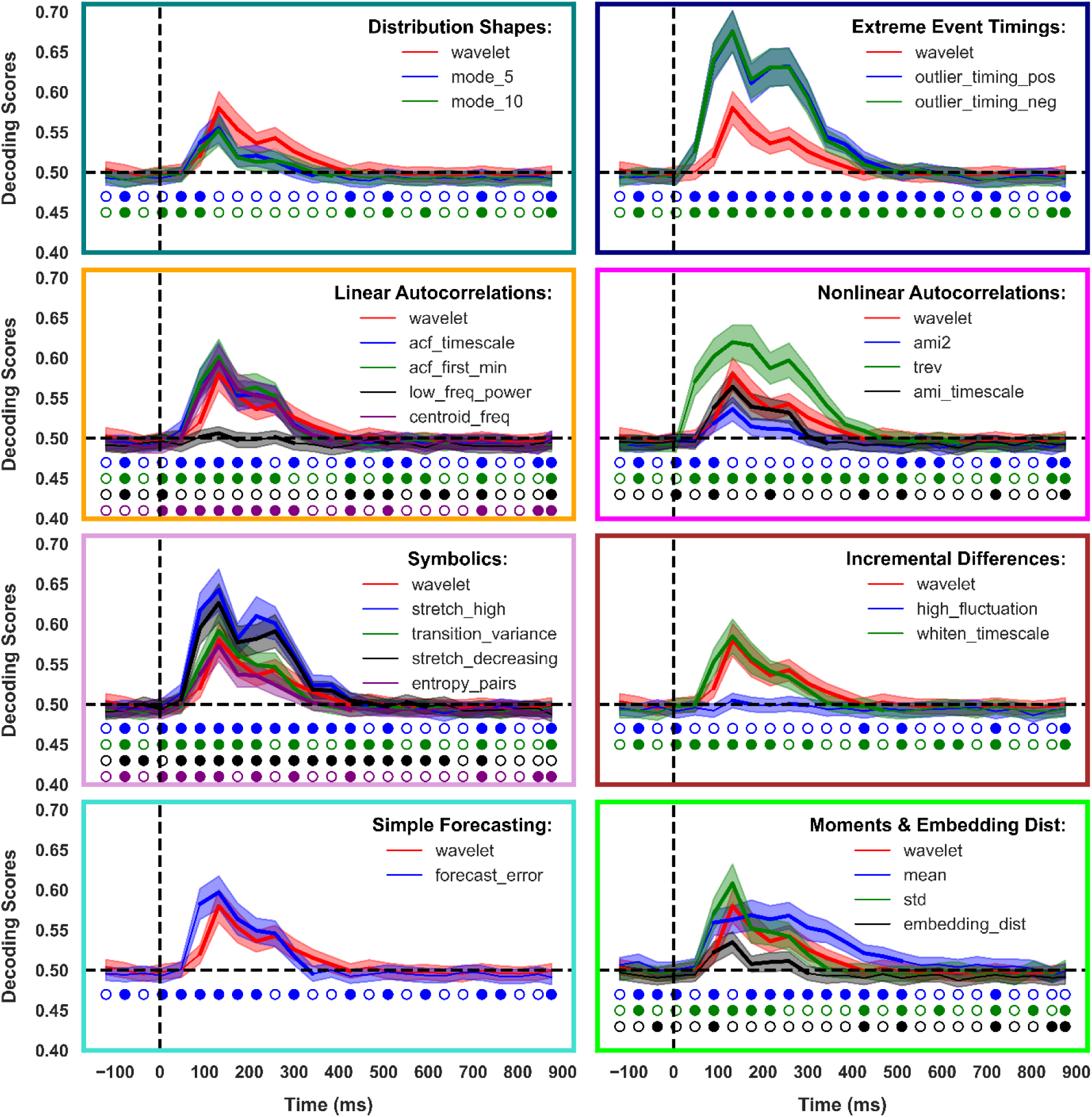
Within-subject decoding of spatial frequencies. Each panel shows the decoding for one category of features over time around the time of stimulus onset (note the distinct boundary colors for each category). Stimuli were presented for 100ms. The dashed vertical line indicates the stimulus onset time, and the dashed horizontal line indicates chance level decoding (0.5). Thickened lines indicate the time points with evidence (BF > 6) for above-chance decoding. Shadings indicate SEM over subjects. Circles indicate the results of Bayes factor comparison between the corresponding decoding and the wavelet decoding results. Filled circles indicate the time point when Bayes factor of the corresponding decoding was higher than the wavelet-based decoding (BF > 6).

The decoding curves followed the established profile of decoding from previous studies. Specifically, the decoding of spatial frequencies showed a significant increase in information coding before 100ms followed by one or two peaks before 400ms before declining back to the baseline (Karimi-Rouzbahani et al., 2021; Karimi-Rouzbahani, 2024).

While many features followed this pattern, there were drastic variations in their temporal profiles including the peak amplitudes and the duration of the above-chance decoding (Figure 3). This suggests that features differ in what aspects of neural activity they capture. For example, the two features of *outlier_timing_pos* and *outlier_timing_neg*, which were within the *extreme event timing* category of features, provided higher decoding values than features of *high_fluctuation* and *whiten_timescale* within the *incremental differences* category of features. The dominance of *extreme event timing* features suggests that, during the first few hundred milliseconds of stimulus presentation, information about different stimuli was reflected in the timings of rapid extreme changes rather than sustained trends in the signals.

When comparing the results of different features against the *wavelet* feature, which was top-performing in previous EEG decoding studies (Taghizadeh-Sarabi et al., 2015; Karimi-Rouzbahani et al., 2021), we saw that some features, but not all, outperformed the *wavelet* feature. For example, *extreme event timing* features provided higher decoding accuracies than the *wavelet* feature in the first 400ms after stimulus onset whereas features within the *distribution shapes* categories could not outperform the *wavelet* feature. This suggests that information might be encoded in different aspects of signals in MEG than EEG.

To quantitatively compare the timing and magnitude of decoding results presented in Figure 3, we extracted three main quantities from the time-resolved decoding curves: average and maximum decoding values and the timing of the maximum decoding in the post-stimulus window (Figure 4). Results confirmed that the features of *extreme event timing* showed higher maximum and average decoding values compared to the rest of features and the *incremental* features were among the worst-performing features (BF > 6: see Bayes factor evidence for pair-wise comparisons between all features in Supplementary Figure 3). While there was insufficient evidence (1/6 < BF <6) for an earlier peak for any of the features, *low frequency power*, *high fluctuation* and *mean* features showed the latest peaks among features (BF > 6).

**Figure 4.**
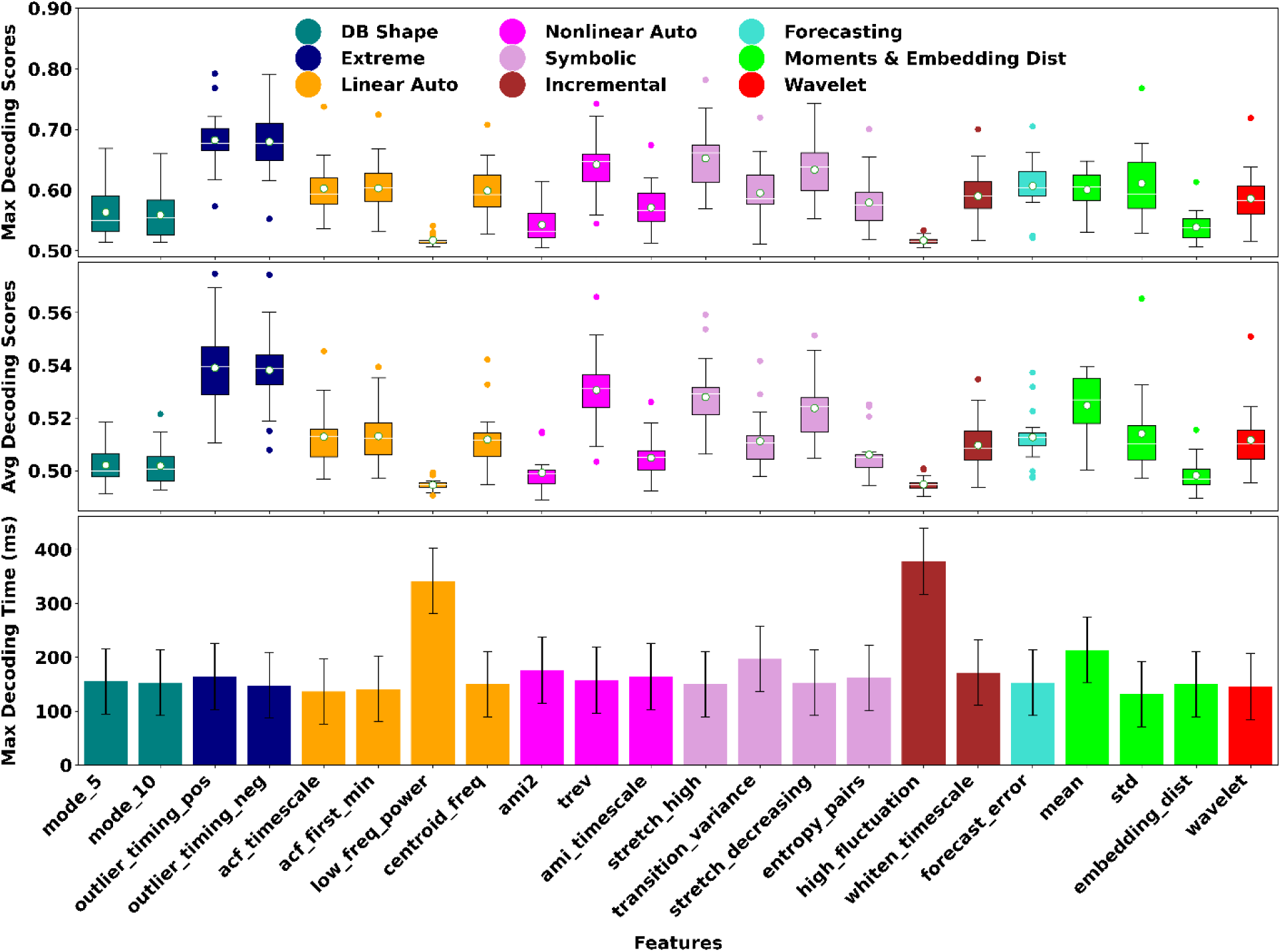
Decoding parameters extracted from *within-subject* decoding results. Panels from top show the maximum decoding, average decoding and the time of maximum decoding for each feature (colors indicate the feature category and correspond to boundary colors used in Figure 3). Box plots show the distribution of data, its quartiles and median and whiskers indicate the maximum and minimum of the data over subjects. Dots indicate outlier data. Bar plots indicate the mean and SEM of the data over subjects.

The decoding of orientations also showed similar rankings of features to spatial frequency, with lower levels of decoding on average (Supplementary Figure 2), which was expected based on previous studies (Quentin et al., 2019; Karimi-Rouzbahani, 2024). The quantification of orientation decoding curves also showed the dominance of *extreme event timing* features (Supplementary Figure 3) and a delayed peak for the *high fluctuation* feature.

Together, these results suggest that the neural codes (i.e., information) were reflected in specific, but not all features of MEG signals. Therefore, extracting the appropriate features of neural activity led to more effective information decoding.

### Cross-Subject Analysis

The above results showed that features related to the timing of extreme or rapid activities provided more information than other features of neural activity within each subject. However, it remains unclear if the same features would capture the neural codes from all subjects. In other words, which features of neural activity are shared by subjects and at what time? Answering this question is not only important for identifying shared neural coding mechanisms across brains but can facilitate brain-computer interface applications. To that end, we used a leave-one-subject-out (LOSO) cross-validation scheme where we trained the classifiers using data from all-minus-one subjects and tested them using the data from an out-of-sample subject.

Similarly to *within-subject* decoding, there were large variations in the decoding profiles across features, including the peak amplitudes and the duration of the above-chance decoding (Figure 5). For example, the two features of *outlier_timing_pos* and *outlier_timing_neg*, which belonged to the *extreme event timings* category of features, provided higher decoding values than features of *high_fluctuation* and *whiten_timescale* within the *incremental differences* category of features. While *cross-subject* decoding profiles showed similarities to the *within-subject* data, there were at least two critical differences as well (Figure 5). First, the level of decoding dropped almost across all features compared to the *within-subject* results (c.f., Figure 3). Second, the temporal profile has slightly changed for some features. For example, many of the features (except for *trev*) showed one peak in *cross-subject* decoding as opposed to the two peaks observed for the *within-subject* analysis. This can mean that the first decoding peak which happened before 200ms, but not the second one which happened after 200ms (c.f., Figure 3), was shared by most subjects. The timing of the generalizable first peak aligns with the well-established dominant sweep of feed-forward flow of visual signals in the brain (Thorpe et al., 1996; Kietzmann et al., 2019).

**Figure 5.**
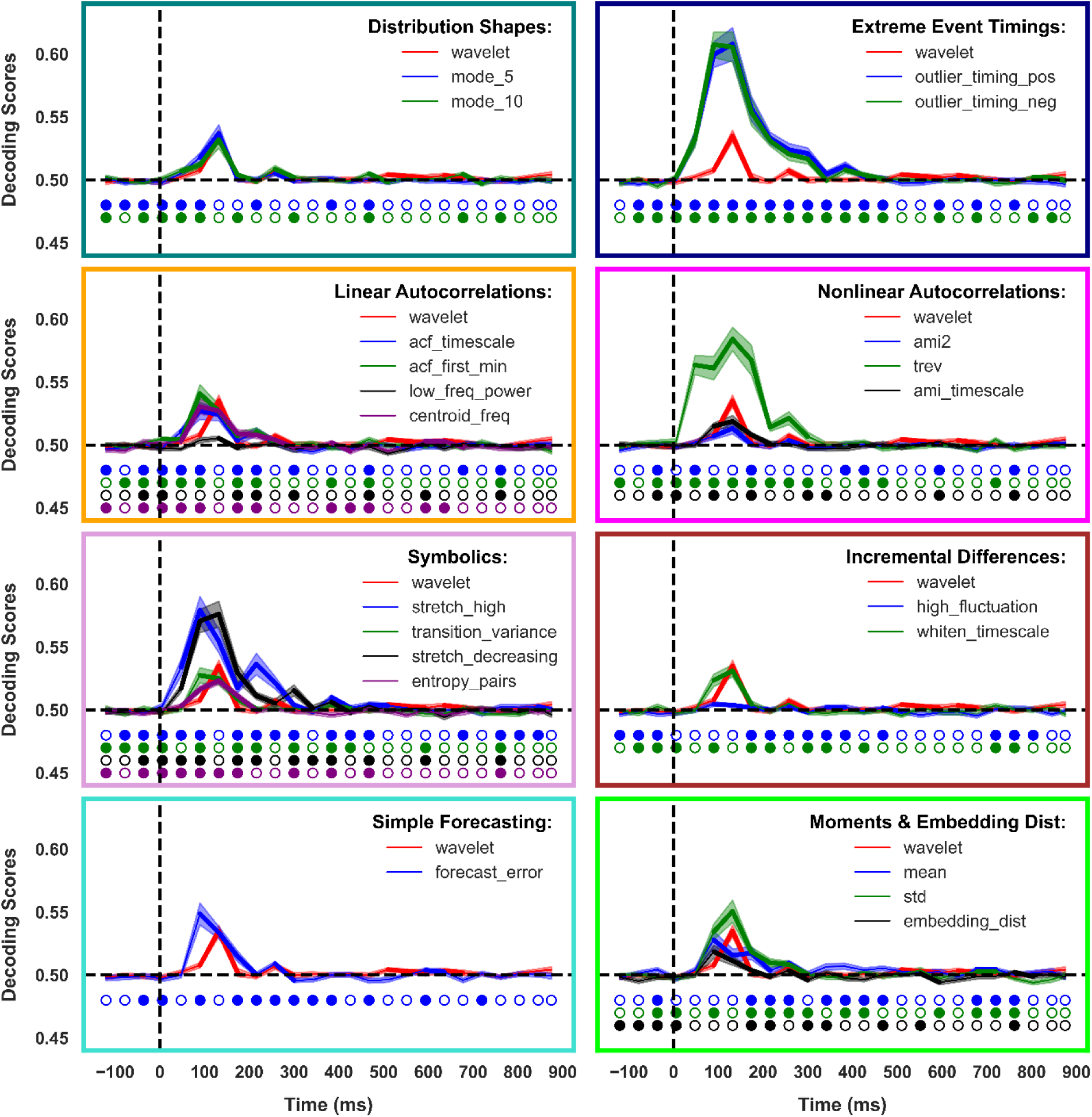
Cross-subject decoding of spatial frequencies. Each panel shows the decoding for one category of features over time around the time of stimulus onset (note the distinct boundary colors for each category). Stimuli were presented for 100ms. The dashed vertical line indicates the stimulus onset time, and the dashed horizontal line indicates chance level decoding (0.5). Thickened lines indicate the time points with evidence (BF > 6) for above-chance decoding. Shadings indicate SEM over subjects. Circles indicate the results of Bayes factor comparison between the corresponding decoding and the wavelet decoding results. Filled circles indicate the time point when Bayes factor of the corresponding decoding was higher than the wavelet-based decoding (BF > 6).

The extracted average and maximum of decoding scores confirmed that the features within *extreme event timing* category showed higher maximum and average decoding accuracies compared to the rest of features and *incremental* features were among the worst-performing features (Figure 6 and BF > 6 in Supplementary Figure 5). We also observed that the timing of the maximum decoding was earlier in *extreme event timing* category than other features, while the *incremental* features showed the most delayed peaks.

**Figure 6.**
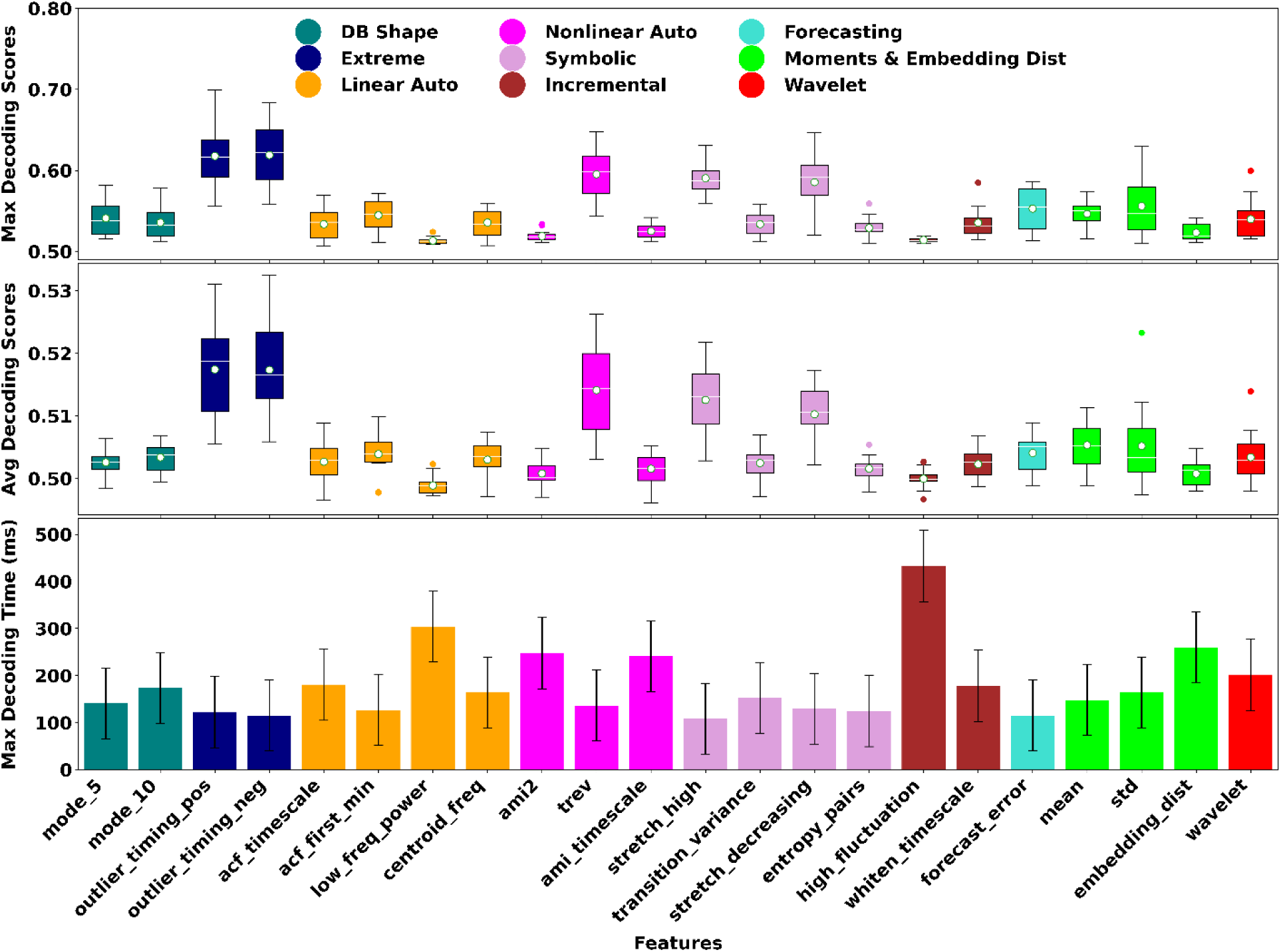
Decoding parameters extracted from cross-subject decoding results. Panels from top show the maximum decoding, average decoding and the time of maximum decoding for each feature (colors indicate the feature category and correspond to boundary colors used in Figure 5). Box plots show the distribution of data, its quartiles and median and whiskers indicate the maximum and minimum of the data over subjects. Dots indicate outlier data. Bar plots indicate the mean and SEM of the data over subjects The *cross-subject* decoding of orientations showed similar results to the spatial frequency (Supplementary Figures 6-8).

### More Informative Features Generalized Better

The above results suggested some consistency between the informative features in *within-* and *cross-subject* analyses. To quantitatively assess this potential consistency we calculated the correlation between the *within-* and *cross-subject* decoding scores. Including all decoding scores (i.e., subjects, features and time windows) in the correlation analysis, we observed a significant positive correlation between the *within-* and *cross-subject* decoding scores (Supplementary Figure 9). Next, to specifically check if this correlation existed for all subjects, we calculated the correlation separately for each subject. This was done for the maximum and average of the decoding curves not to be affected by the similar temporal dynamics of the decoding in the *within-* and *cross-subject* analyses. Consistently across subjects, the correlations were positive (and significant p < 0.01; except for subjects 1, 2 and 8 in average decoding) between the level of decoding in the *within-subject* and *cross-subject* analyses across features (Figure 7). This suggests that the features which were most effective in the *within-subject* decoding were also most effective in the *cross-subject* decoding consistently across subjects. We observed dominantly positive patterns of correlation for the orientation decoding results as well (Supplementary Figure 10).

**Figure 7.**
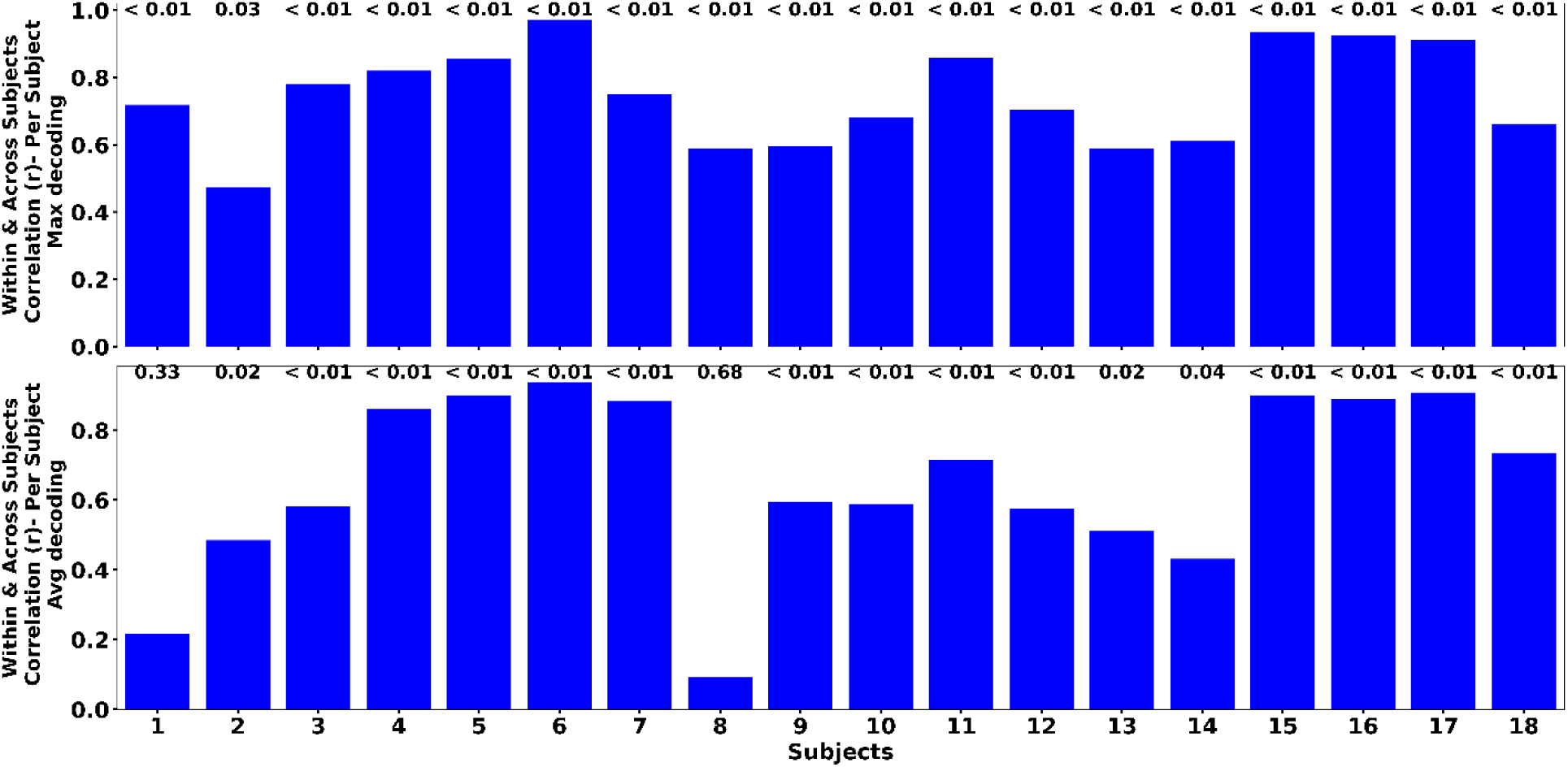
Correlation between the decoding parameters of within- and cross-subject decoding analyses of spatial frequency. Pearson correlation was calculated using the results separately for each subject and over features. Numbers above the bars indicate correlation p-values.

Together these results confirm that distinct set of features allowed for different levels of decoding and cross-subject generalizability in MEG. Moreover, these suggest that features which enabled more effective stimulus decoding also facilitated *cross-subject* generalizability.

### Shared and Subject-Specific Neural Codes

The above *cross-subject* results suggested that earlier time windows of signals (i.e., 0-200ms) seemed more generalizable across subjects than the later time windows (i.e., after 200ms; Figure 5). We also found support for this by comparing the timing of the maximum decoding which happened earlier in the *cross-* than *within-subject* decoding (BF = 189; Figure 8A). To test if *cross-subject* generalizable decoding mainly relied on earlier than later neural codes, we calculated the percentage of above-chance (BF > 6) decoding in the time window before and after the 200ms mark. We saw a higher percentage of the above-chance decoding in the *cross-* than *within-subject* decoding in the time window before 200ms (BF = 42; Figure 8B). This suggests that *cross-subject* generalizable neural codes were more common in earlier than later (i.e., after 200ms) time windows. There was also a higher percentage of above-chance decoding for the *cross-* than *within-subject* decoding in the time window before 200ms (BF = 95; Supplementary Figure 11).

**Figure 8.**
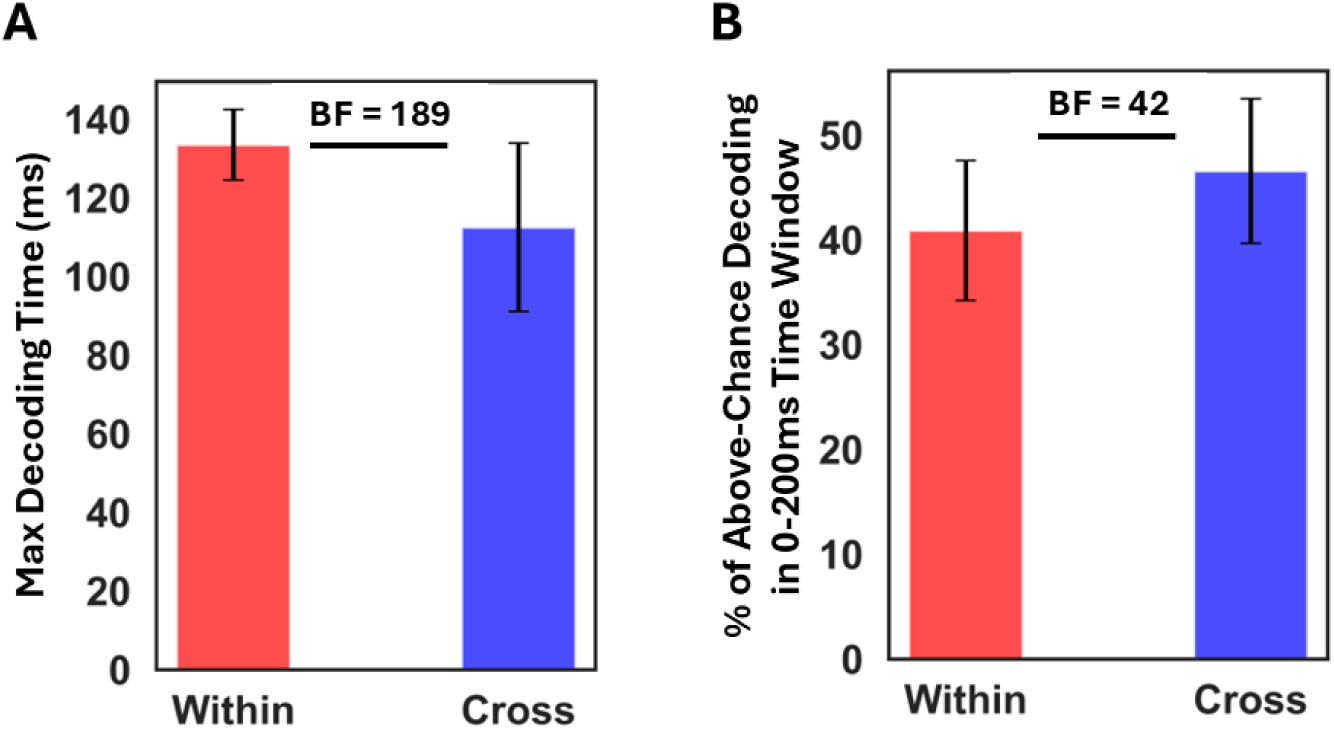
Comparison of the timing of spatial frequency decoding in *within-* and *cross-subject* decoding. **A** shows the time of maximum decoding in the within- and cross-subject decoding analyses. **B** shows the percentage of above-chance decoding scores found in the earlier time window (0-200ms) in the within- and cross-subject decoding analyses. Error bars indicate SEM over subjects.

These results suggested that patterns of informative neural codes might be more consistent across subjects in the earlier than later windows of evoked activity. To test this hypothesis more directly, we evaluated which features provided the maximum decoding in the earlier (before 200ms) and later (after 200ms) time windows separately for each subject (Figure 9). If neural codes were more consistent across subjects in the earlier than later windows, we should see dominance of one feature over the others in the *cross-subject* decoding results in earlier but a more distributed set of informative features in the later time windows. As predicted, in the earlier window of 0-200ms, we observed the dominance of two features from *extreme event timings* which provided the highest decoding for 13 out of 18 subjects (Figure 9). In the later time window (after 200ms) a higher number of features were informative with *extreme event timings* being dominated by *trev* and *stretch_ligh*. The *mode_5* and *mean* features were also informative in later but not earlier time window. These results suggest that while the informative neural codes were more consistent across the brains in earlier time windows (0-200ms), there were more distinct patterns of informative neural codes across individuals in later time windows (200ms onwards). This effect was even more pronounced in orientation decoding (Supplementary Figure 12).

**Figure 9.**
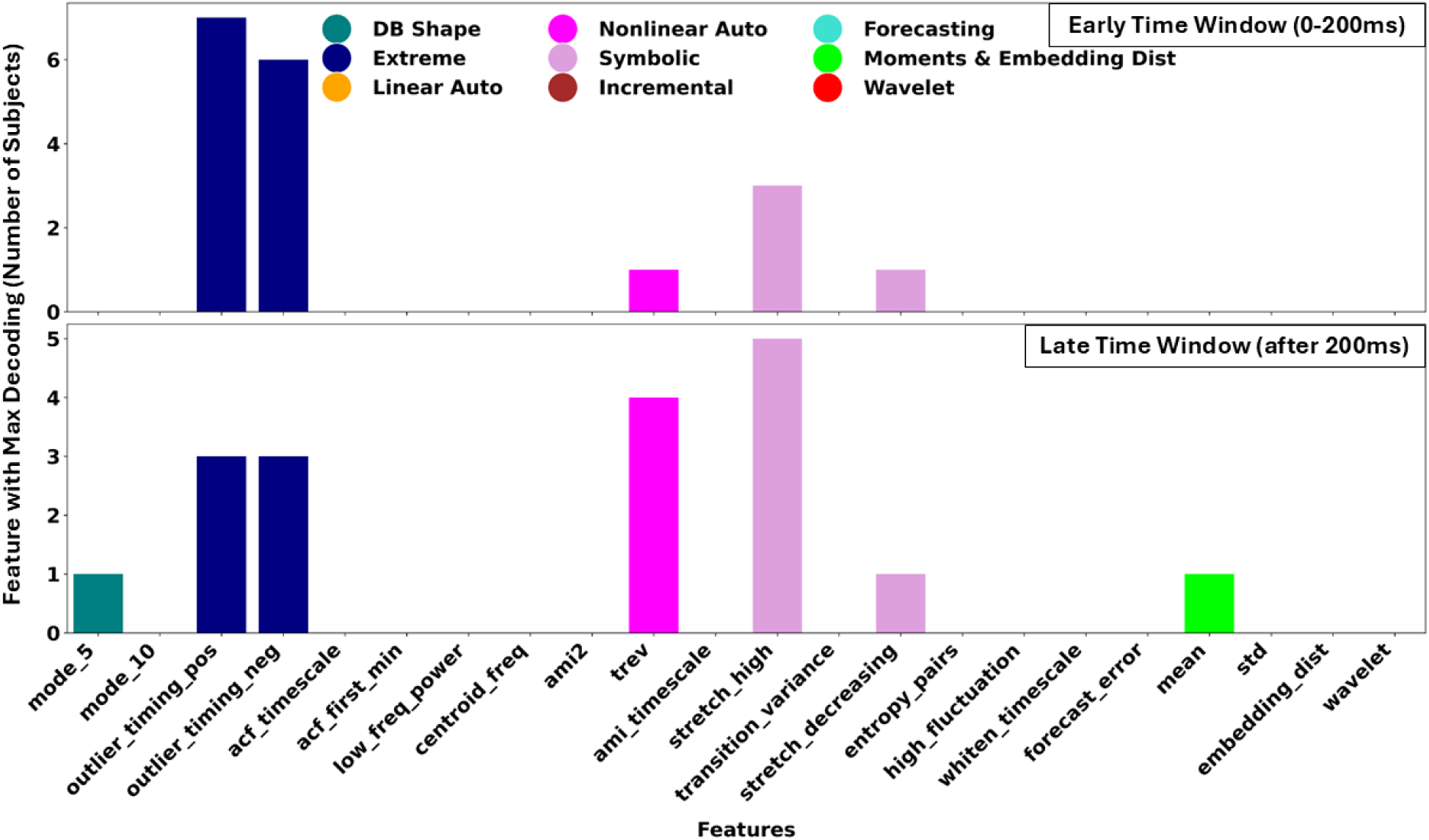
Distribution of features which provided the highest spatial frequency decoding. Top and bottom panels show the results for the earlier (0-200ms) and later (after 200ms) analysis window, respectively.

### Relationship to Behavior

The above results offer insights into the features which enabled more effective decoding and generalizability across subjects. However, it remained unclear if the brain also used such information coding mechanisms for cognitive processing (i.e., visual perception here). One way to validate whether a feature of neural activity is used by the brain or not is to see if it can predict behavioral performance in the same experiment (Ritchie et al., 2015; Karimi-Rouzbahani et al., 2019; Karimi-Rouzbahani et al., 2021). To check if an enhancement in decoding corresponded to improved prediction of behavior, we evaluated the correlation between the decoding and the subjects’ average behavioral reaction times (Karimi-Rouzbahani et al., 2021) and then evaluated the correlation of the resultant values against the decoding scores of the same subjects. A negative correlation would mean that enhanced decoding, as a result of more effective feature extraction, enhanced behavioral prediction (Ritchie et al., 2015; Karimi-Rouzbahani et al., 2021). However, neither maximum nor average decoding accuracy showed a significant (p < 0.01) level of negative correlation to behavioral performance (Figure 10). This non-significant correlation can be explained by the complex nature of the experimental task involving more components than just a simple stimulus discrimination.

**Figure 10.**
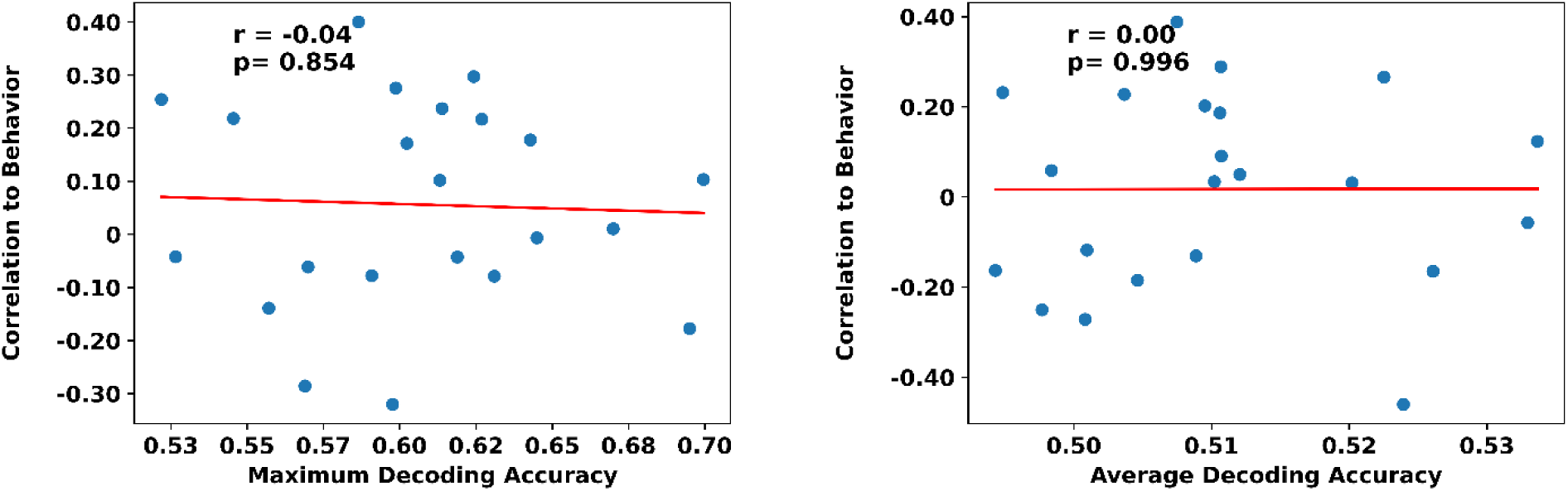
Correlation between the parameters of *within-subject* decoding (maximum: left; average: right) and the behavioral reaction times for spatial frequency. Pearson correlation was calculated between the decoding and reaction time followed by a second-level correlation between the results and the decoding results. Each dot represents data from one feature. The slant line shows the best linear fit to the data.

## Discussion

The primary objective of this study was to identify *within-subject* and *cross-subject* generalizable features of neural activity in MEG decoding. We extracted 22 of the most informative features of neural time series from our MEG dataset in a time-resolved decoding pipeline to discriminate spatial frequencies and orientations of grating stimuli. Results showed more robust decoding of gratings’ spatial frequency than orientation. Importantly, we found that specific features of neural activity, which reflected rapid and drastic changes in signal amplitudes, achieved higher decoding accuracy consistently across subjects, particularly within the initial time window of stimulus presentation (i.e., before 200 ms). Importantly, the same features generalized across subjects most robustly, suggesting that sharp changes in brain activity were key sources of information for decoding. The analysis of interpretable features of neural activity not only provides new insights into the potential neural coding mechanisms implemented by the brain but also provides a more explainable approach for real-world BCI applications.

### Shared and Subject-Specific Neural Codes

One main question of this study was what do the brains share about information coding? Is it the timings of information coding, the patterns of neural activity that carries the information or a combination of both? Our results showed that different brains shared specific patterns of activity within specific time windows. More specifically, we found that the most *cross-subject* generalizable patterns of activity were rapid, temporary, and extreme signal deviations (as captured by *extreme event timing* features) which occurred early after the stimulus onset (i.e., 100-200ms; Figure 5). In terms of timing, the appearance of stimulus information shortly after stimulus appearance aligns well with established literature on visual evoked responses. This time frame supports the early stages of visual processing in the visual cortex, where rapid neural changes occur in response to stimuli and are assumed to be relatively consistent across subjects (Thorpe et al., 1996; Kietzmann et al., 2019). In terms of patterns, the extreme events in the signal capitalize on the high signal-to-noise ratio (SNR) after the stimulus onset by isolating the most significant deviations, effectively filtering out less informative (and potentially subject-specific) signal variations, leading to higher generalizations early after the stimulus onset.

We observed that while the *within-subject* decoding curves remained above-chance for a prolonged period after this initial sweep (Figure 3), the *cross-subject* generalization decoding curves dropped shortly after this initial sweep (Figure 5). This suggested that most brains produced more similar neural codes immediately after the stimulus onset which changed to more subject-specific patterns after the initial sweep of signals (Figure 9). While the initial sweep of information is dominantly time-locked to the stimulus onset, leading to more consistency across the brains, as time passes, the patterns become more individualized as subjects implement their personal mental processes and strategies to perform the task. We supported this by showing a higher number of distinct features contributing to decoding after than before the 200ms mark (Figure 9 and Supplementary Figure 11). This is also supported by our recent study showing that the timescale of neural codes varies over the course of the trial, with longer neural codes in the initial phase of stimulus presentation, followed by shorter neural codes in the following windows (Karimi-Rouzbahani, 2024).

### Neural Codes in MEG vs EEG

Many of the features extracted from our MEG dataset outperformed the *wavelet* feature which had been the top-performing feature in previous EEG studies (Taghizadeh-Sarabi et al., 2015; Karimi-Rouzbahani et al., 2021). This can be explained by several factors, including the distinct nature of MEG vs EEG. While both modalities provide signals with high temporal resolution, MEG is more sensitive to tangential sources vs EEG which is sensitive to more radial sources. This can lead to distinct sources of information captured by the two modalities (Gross, 2019). Moreover, while in EEG we evaluated all the informative features from previous studies, here our 22 features had been selected from a large set of >7,700 features (Lubba et al., 2019), potentially enhancing the variance that we captured with our MEG features. Finally, the set of visual stimuli and conditions which were decoded in previous studies were semantic object categories (Taghizadeh-Sarabi et al., 2015; Karimi-Rouzbahani et al., 2021) rather than the low-level grating stimuli used here, which might be encoded differently in the brain. Our future goal is to systematically compare information coding in EEG and MEG using similar datasets.

Neural codes can be defined both across time and space (Panzeri et al., 2010). Knowing the poor spatial resolution of MEG (and EEG) compared to fMRI, here we extracted features from each MEG channel separately before concatenating all channels for decoding. This focused our study on *temporal* rather than *spatial* neural codes, which is an additional aspect of information coding worth considering in the future using fMRI.

### Relevance of Neural Codes

More effective decoding of neural activity, which can be achieved by extracting more relevant features from the signals, might not necessarily mean better access to the neural codes. In other words, the brain might not use the extracted neural code for its cognitive processes. One way to validate whether a neural code is used by the brain or not is to check if it can predict the behavioral performance. As opposed to several previous studies (Ritchie et al., 2015; Karimi-Rouzbahani et al., 2021; Karimi-Rouzbahani et al., 2019), correlations between decoding accuracy and behavioral performance were non-significant in the present study. This can be explained by the nature of the complex experimental task which involved not only the presentation of stimulus but also cue, memory and stimulus matching processes (Quentin et al., 2019), each of which might have affected the reaction time. In the future, it is interesting to decode each of these aspects separately and try to evaluate the contribution of each aspect in predicting the behavioral performance ideally on a trial-by-trial basis. For that, we will need a dataset with a significantly larger number of trials than the 400 trials collected in the present study for each subject.

### Implications for Brain-Computer Interface Systems (BCIs)

The present study proposes a promising direction for BCI development by focusing on distinct, rapid changes in brain activity observed before 200 milliseconds following a stimulus. These specific and reliable neural patterns offer a compelling complementary neural code to the established neural codes used in BCI systems such as P300 or motor imagery (Aggarwal et al., 2022). Our results suggest that earlier windows of evoked signals, which correspond to the early sweep of visual information in the brain, are robust and *cross-subject* generalizable, which can be utilized in addition to the conventional P300 signals (i.e., a positive deflection in the brain’s electrical activity that appears around 300 milliseconds after a stimulus) which rely on later processing stages of the stimulus (Guger et al., 2009). These neural codes, if utilized, might help to overcome the performance challenges that the current P300-based BCI systems face (Fazel-Rezai et al., 2012; Pan et al., 2022).

A key contribution of this study is its deliberate emphasis on interpretable features for neural decoding, a contrast to the “black-box” methodologies exemplified by deep learning methods (Rajpura et al., 2024). While deep learning has demonstrated success in decoding brain signals, its inherent lack of transparency makes it difficult to ascertain which specific signal characteristics are driving its predictions and their generalizability across different users. This study demonstrates that by prioritizing features that are both interpretable and generalizable – specifically early, transient responses – competitive decoding performance can be achieved. Furthermore, unlike prior studies (e.g., Csaky et al. 2023; Khademi et al., 2023) which employed complex deep learning techniques with limited interpretability, this research systematically evaluates the *cross-subject* applicability of these sharp activity changes, providing a novel framework for future BCI systems.

### Conclusion

This study emphasizes the significant value of systematically evaluating a wide range of interpretable MEG-based features for neural decoding. Utilizing a Random Forest classifier on high-resolution MEG data, the research identified key features that demonstrated high accuracy in both within-subject and, importantly, cross-subject decoding, showing strong generalization across individuals. The findings underscore the advantages of MEG, particularly its sensitivity to rapid temporal changes in brain activity, which proves crucial for decoding visual stimulus and developing more transparent and user-friendly BCIs. The peak decoding accuracy observed within the earlier post-stimulus window highlights this period as particularly informative for neuroscientific applications focused on visual stimuli, potentially offering a pathway to computationally more efficient BCI designs.

Our findings highlight the value of a systematic, feature-based approach to neural decoding, providing a detailed understanding of the informative aspects of MEG signals. This work lays the foundation for future studies aimed at building robust, generalizable, and interpretable BCI systems that can effectively translate neural signals into actionable outputs for real-world applications.

## Acknowledgements

HKR was supported by Australian Research Council’s DECRA Fellowship DE230100608 and Newton International Fellowship follow-on grants AL231037 & AL24100035 from the UK Royal Society.

## Supplementary Materials

**Supplementary Figure 1.**
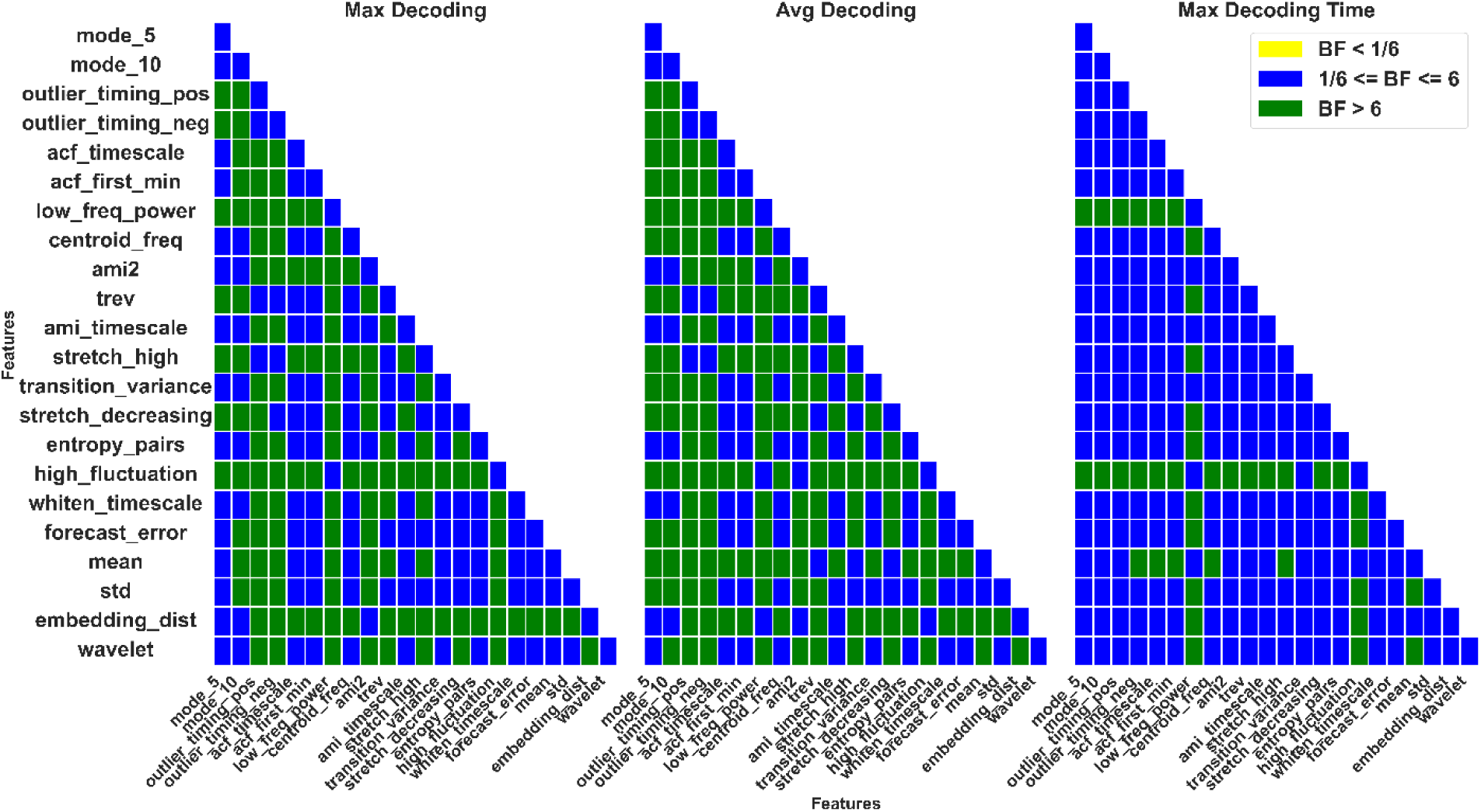
Pair-wise Bayes factor evidence for difference between quantities extracted from *within-subject* spatial frequency decoding curves. Each matrix element indicates color-coded Bayes factor between one pair of features. Yellow indicates evidence smaller than 1/6, blue between 1/6 and 6, and green indicates evidence greater than 6.

**Supplementary Figure 2.**
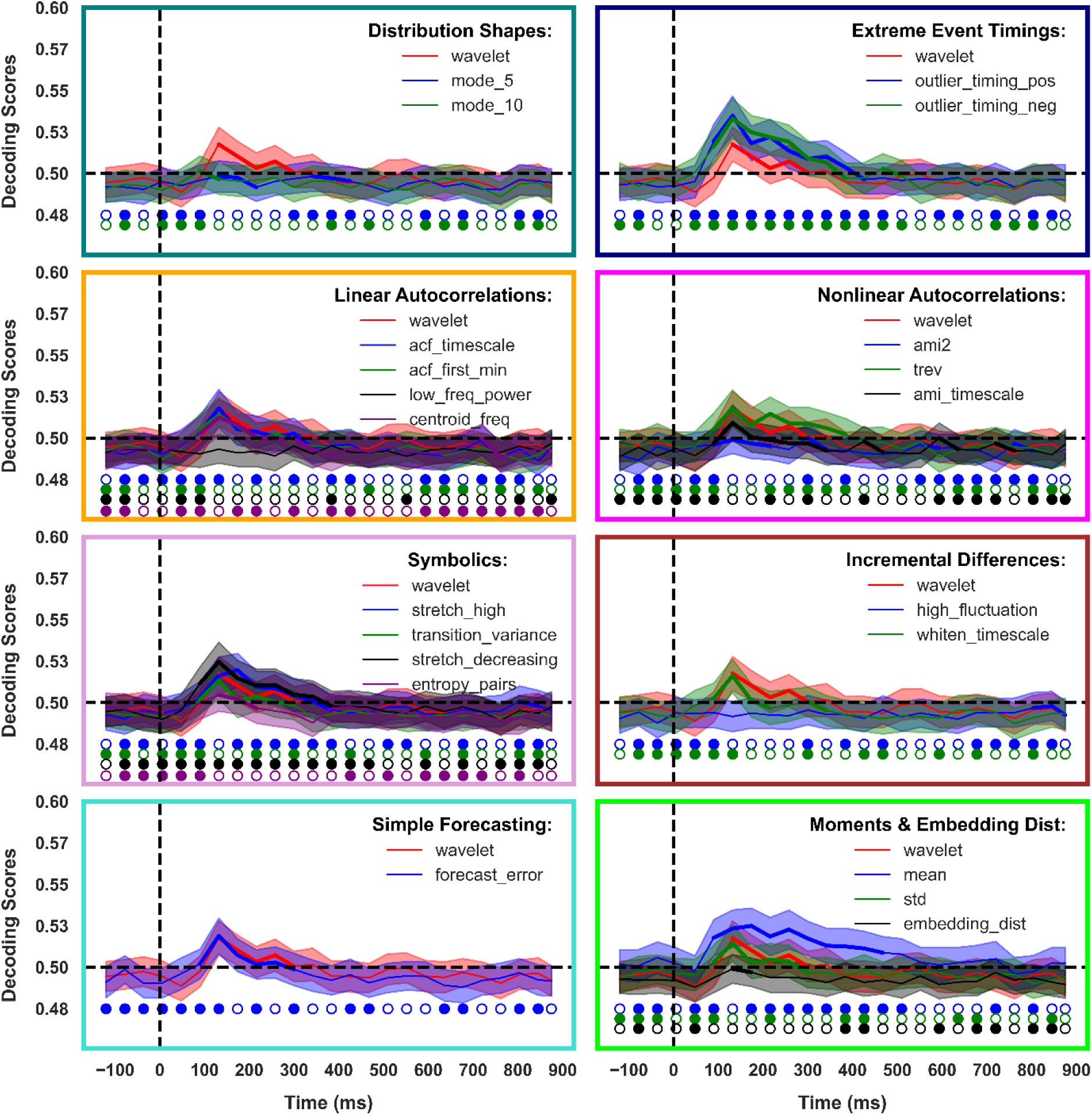
*Within-subject* decoding of orientations. Each panel shows the decoding for one category of features over time around the time of stimulus onset (note the distinct boundary colors for each category). Stimuli were presented for 100ms. The dashed vertical line indicates the stimulus onset time, and the dashed horizontal line indicates chance level decoding (0.5). Thickened lines indicate the time points with evidence (BF > 6) for above-chance decoding. Shadings indicate SEM over subjects. Circles indicate the results of Bayes factor comparison between the corresponding decoding and the wavelet decoding results. Filled circles indicate the time point when Bayes factor of the corresponding decoding was higher than the wavelet-based decoding (BF > 6).

**Supplementary Figure 3.**
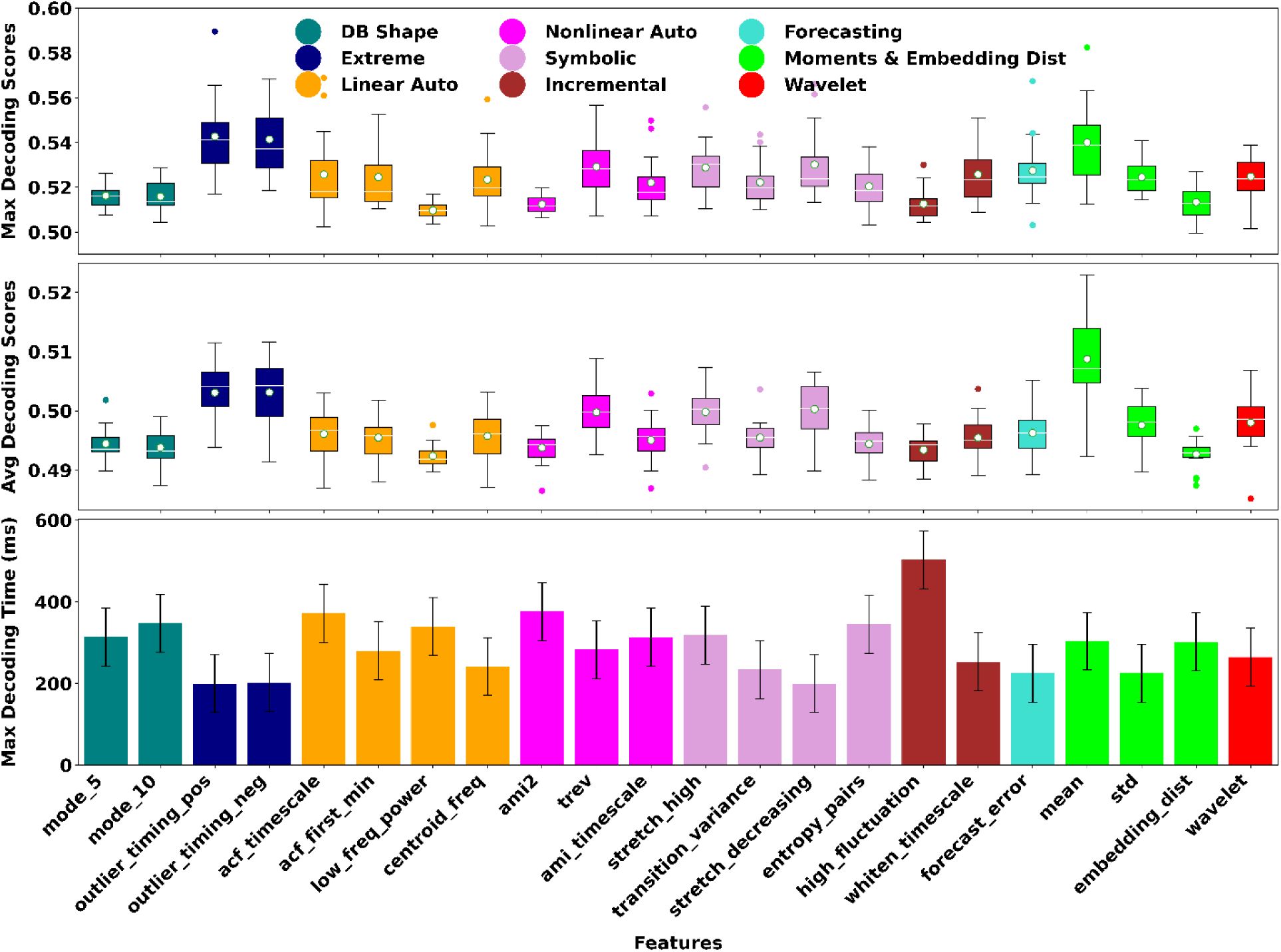
Decoding parameters extracted from *within-subject* decoding analysis. Panels from the top show the maximum decoding, average decoding and the time of maximum decoding for each feature (colors indicate the feature category and correspond to boundary colors used in Supplementary Figure 2). Box plots show the distribution of data, its quartiles and median and whiskers indicate the maximum and minimum of the data over subjects. Dots indicate outlier data. Bar plots indicate mean and SEM of the data over subjects.

**Supplementary Figure 4.**
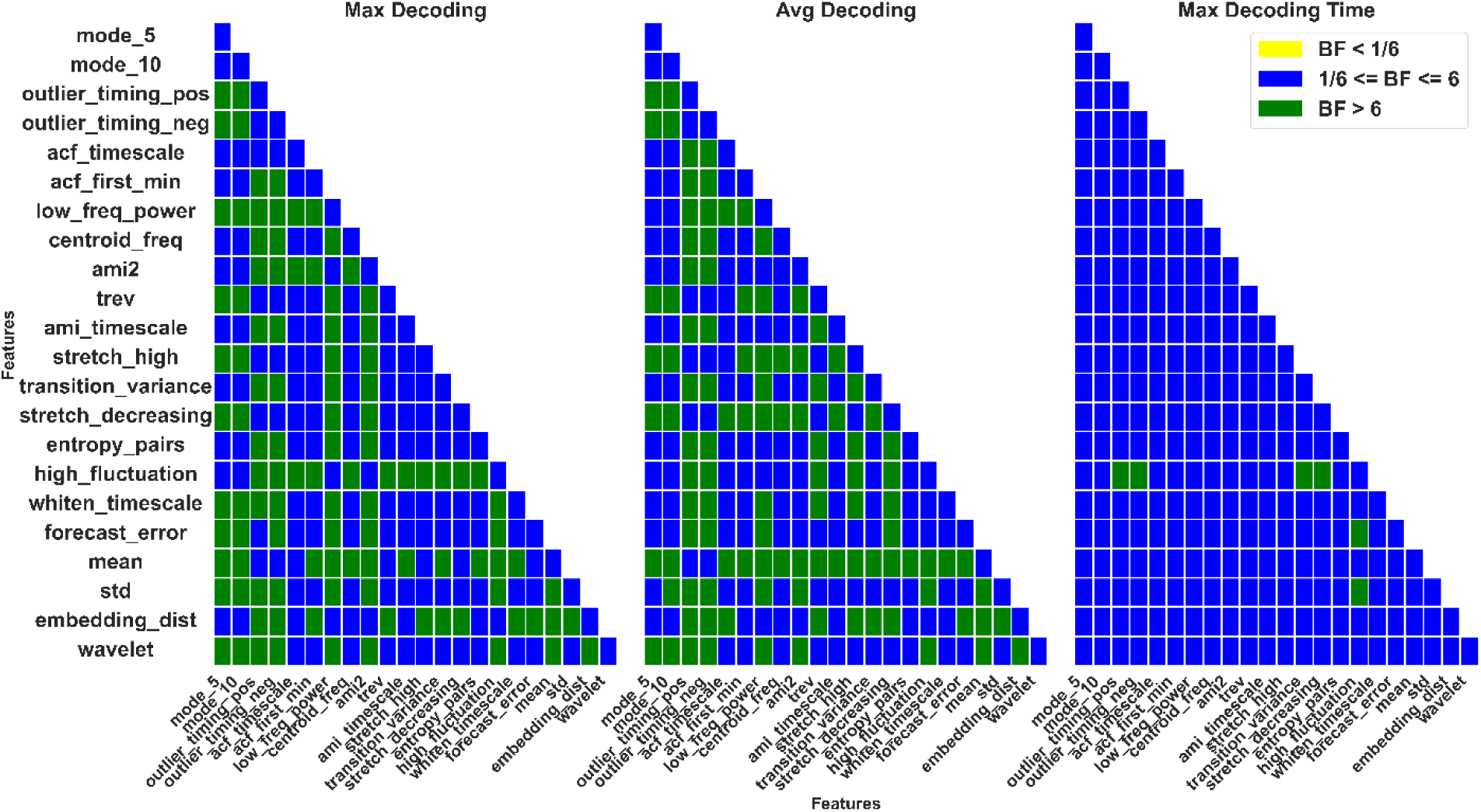
Pair-wise Bayes factor evidence for difference between quantities extracted from *within-subject* orientation decoding curves. Each matrix element indicates color-coded Bayes Factor between one pair of features. Yellow indicates evidence smaller than 1/6, blue between 1/6 and 6, and green indicates evidence greater than 6.

**Supplementary Figure 5.**
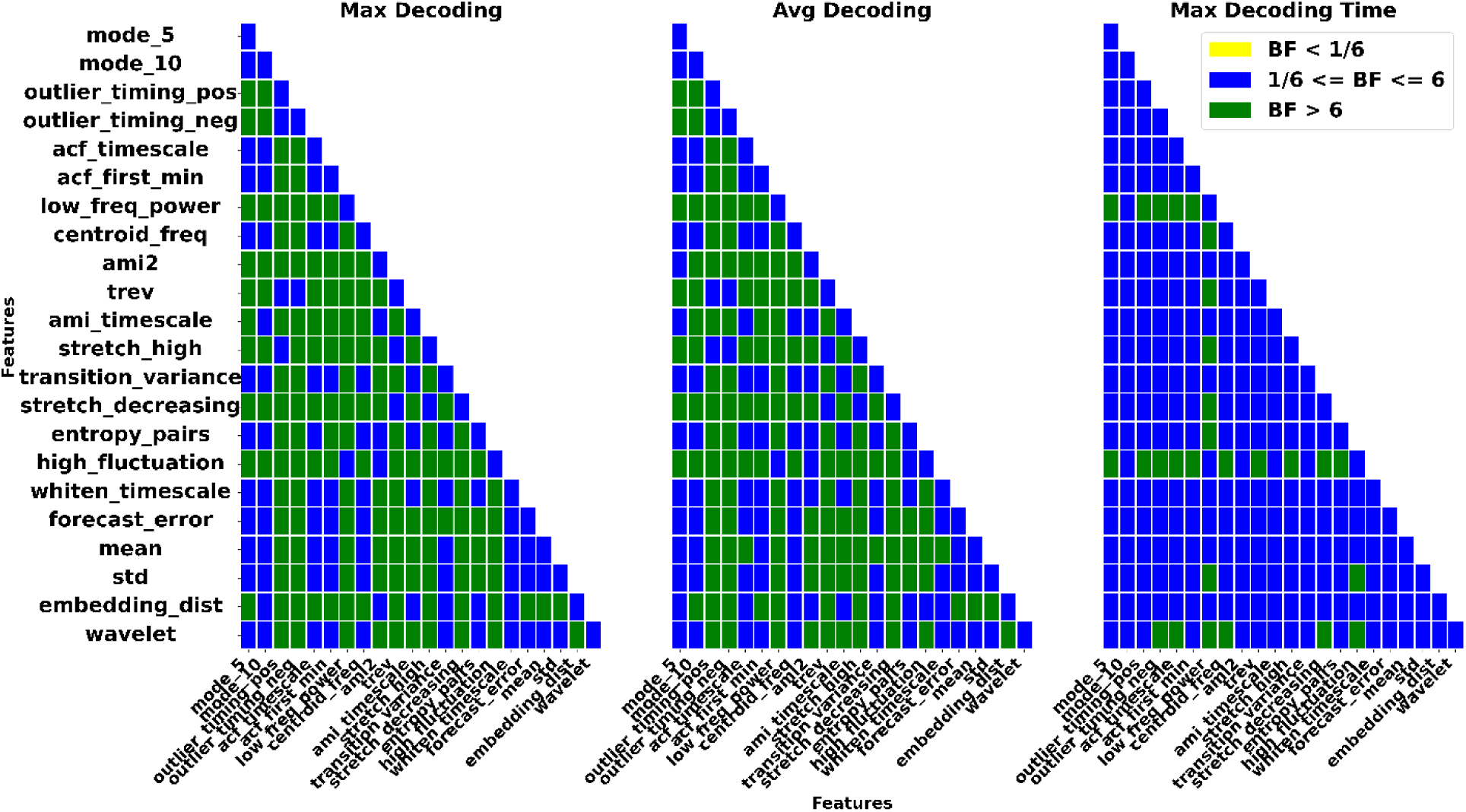
Pair-wise Bayes factor evidence for difference between quantities extracted from *cross-subject* spatial frequency decoding curves. Each matrix element indicates color-coded Bayes factor between one pair of features. Yellow indicates evidence smaller than 1/6, blue between 1/6 and 6, and green indicates evidence greater than 6.

**Supplementary Figure 6.**
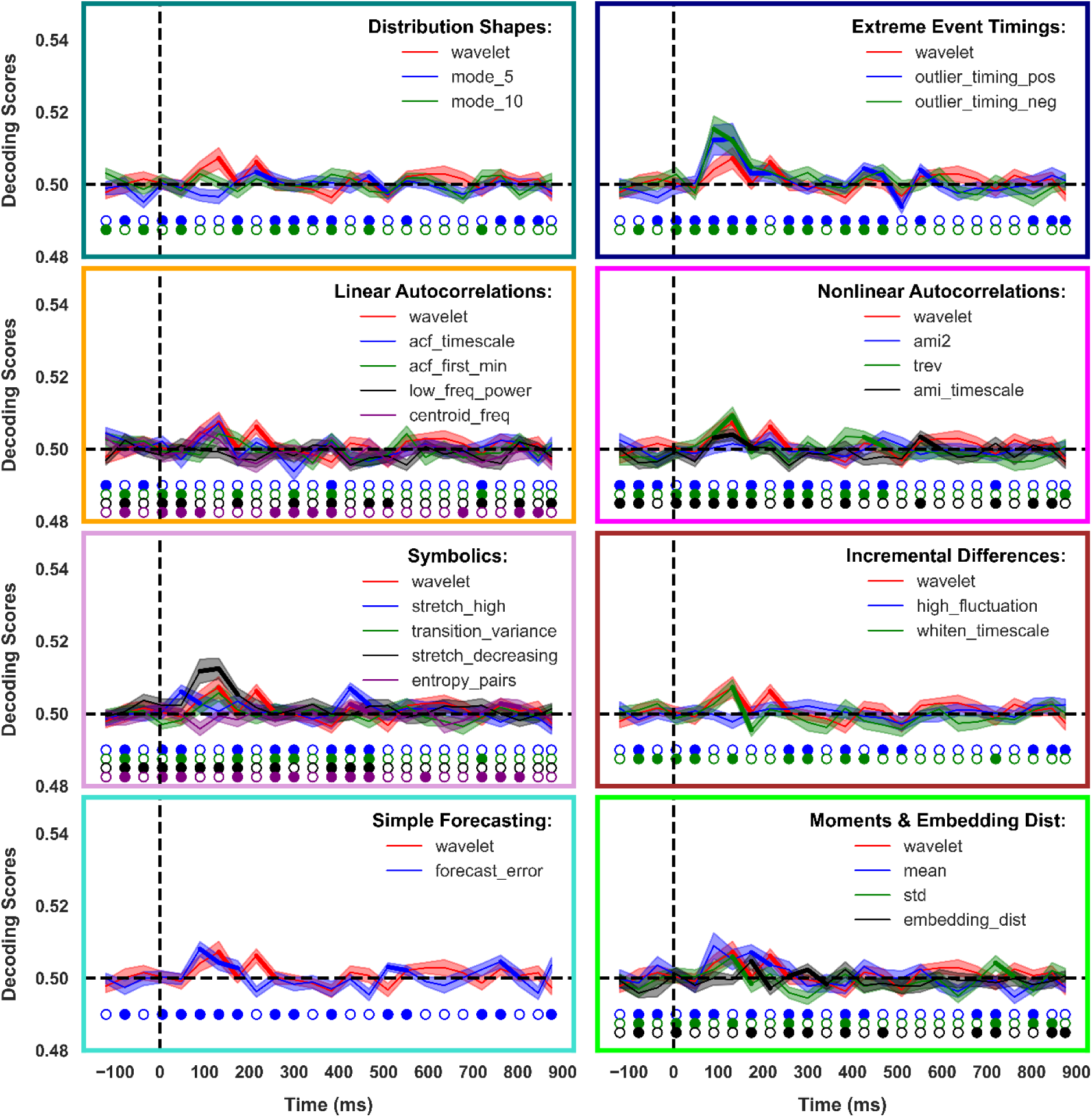
*Cross-subject* decoding of orientation. Each panel shows the decoding for one category of features over time around the time of stimulus onset (note the distinct boundary colors for each category). Stimuli were presented for 100ms. The dashed vertical line indicates the stimulus onset time, and the dashed horizontal line indicates chance level decoding (0.5). Thickened lines indicate the time points with evidence (BF > 6) for above-chance decoding. Shadings indicate SEM over subjects. Circles indicate the results of Bayes factor comparison between the corresponding decoding and the wavelet decoding results. Filled circles indicate the time point when Bayes factor of the corresponding decoding was higher than the wavelet-based decoding (BF > 6).

**Supplementary Figure 7.**
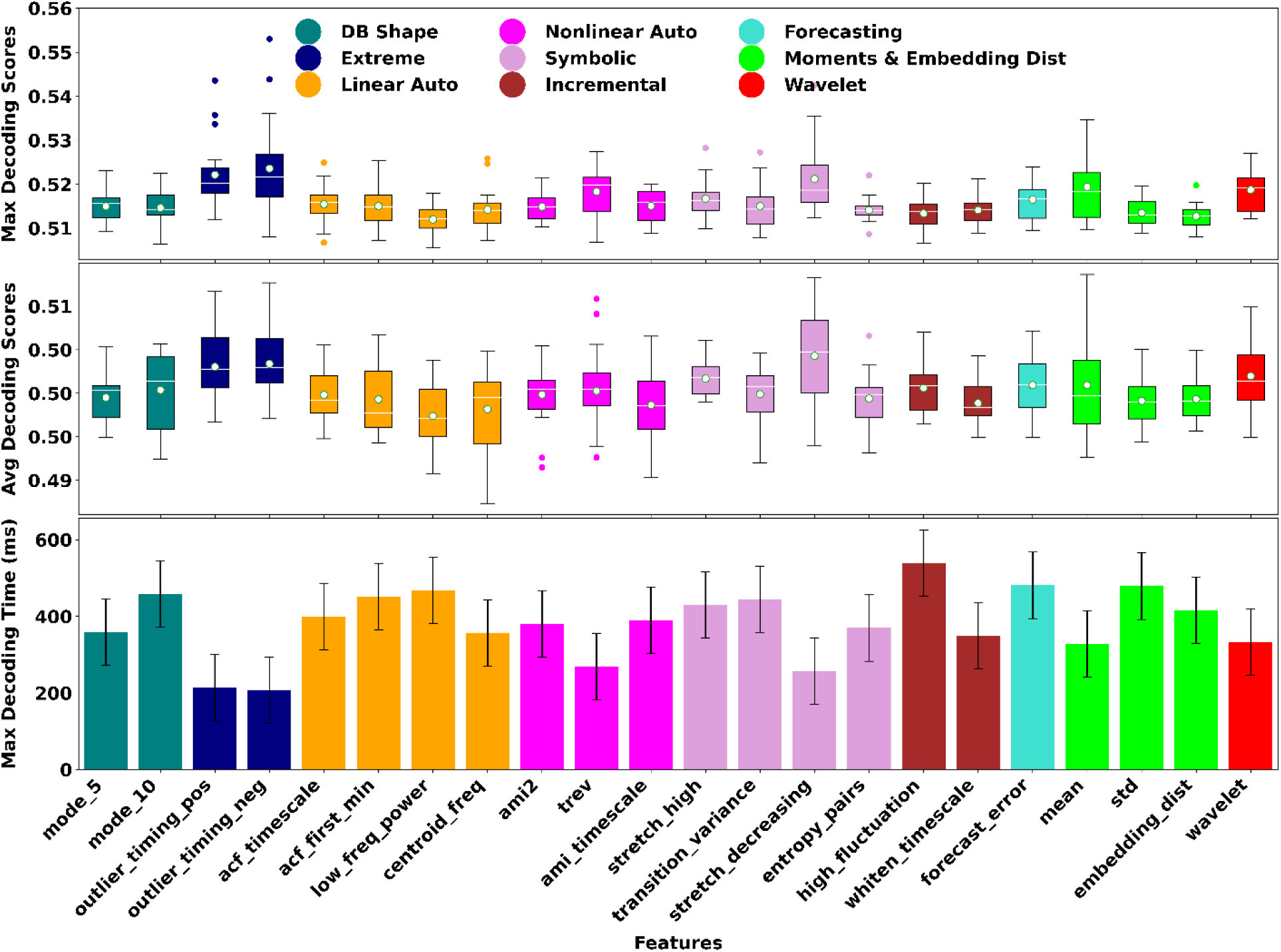
Decoding parameters extracted from *within-subject* decoding analysis. Panels from the top show the maximum decoding, average decoding and the time of maximum decoding for each feature (colors indicate the feature category and correspond to boundary colors used in Supplementary Figure 6). Box plots show the distribution of data, its quartiles and median and whiskers indicate the maximum and minimum of the data across subjects. Dots indicate outlier data. Bar plots indicate mean and SEM of the data across subjects.

**Supplementary Figure 8.**
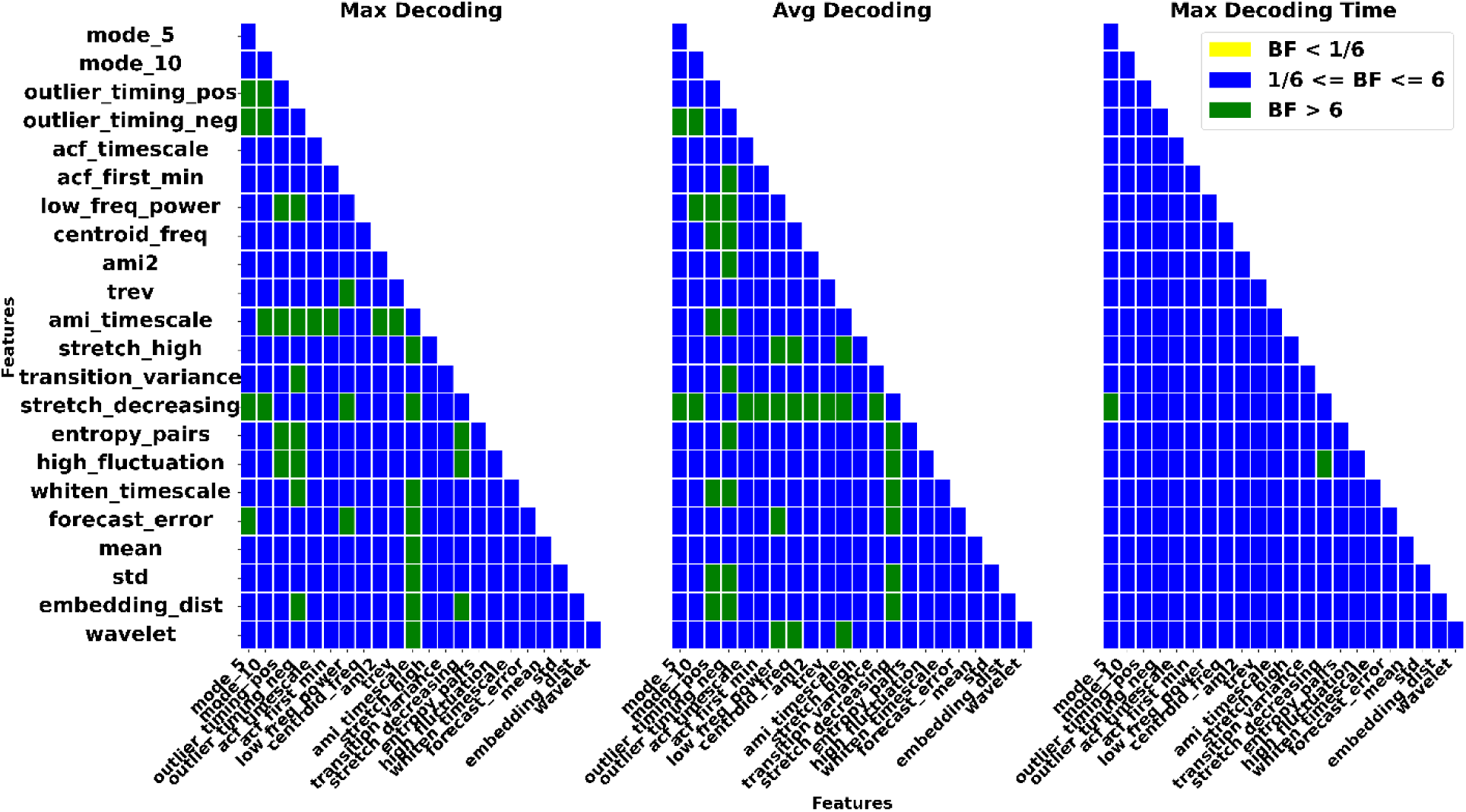
Pair-wise Bayes factor evidence for difference between quantities extracted from *cross-subject* orientation decoding curves. Each matrix element indicates color-coded Bayes factor between one pair of features. Yellow indicates evidence smaller than 1/6, blue between 1/6 and 6, and green indicates evidence greater than 6.

**Supplementary Figure 9.**
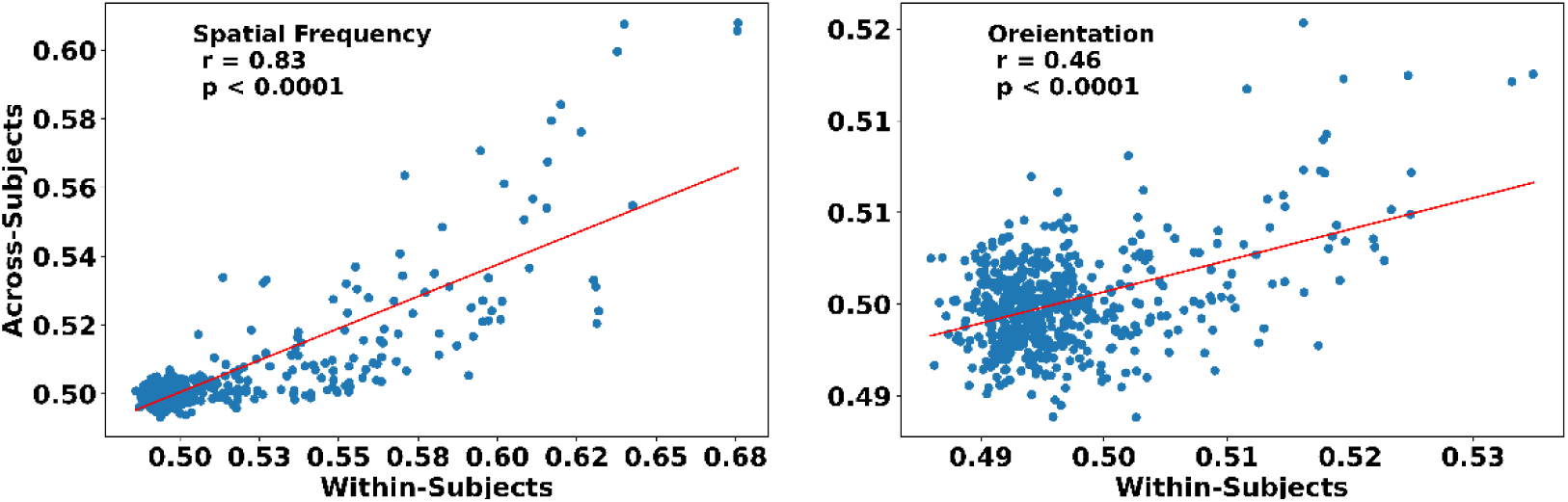
Correlation between the decoding results of *within-* and *cross-subject* decoding analyses of spatial frequency (left) and orientation (right). Pearson correlation was calculated using the combined data from all subjects, features and time windows across the trial (dots). The slant line shows the best linear fit to the data.

**Supplementary Figure 10.**
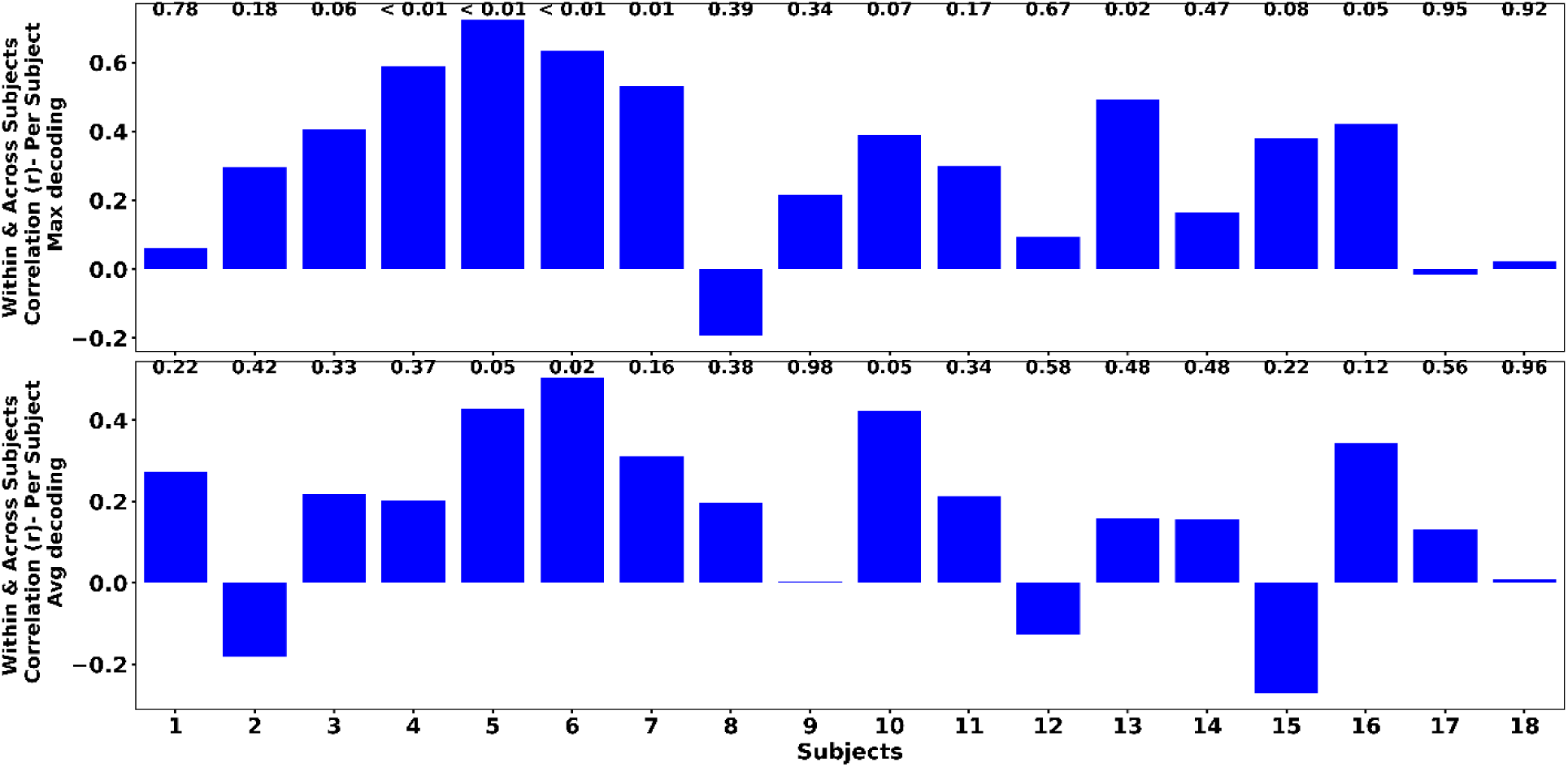
Correlation between the decoding parameters of *within-* and *cross-subject* decoding analyses of spatial frequency. Pearson correlation was calculated using the results separately for each subject and over features. Numbers above the bars indicate correlation p-values.

**Supplementary Figure 11.**
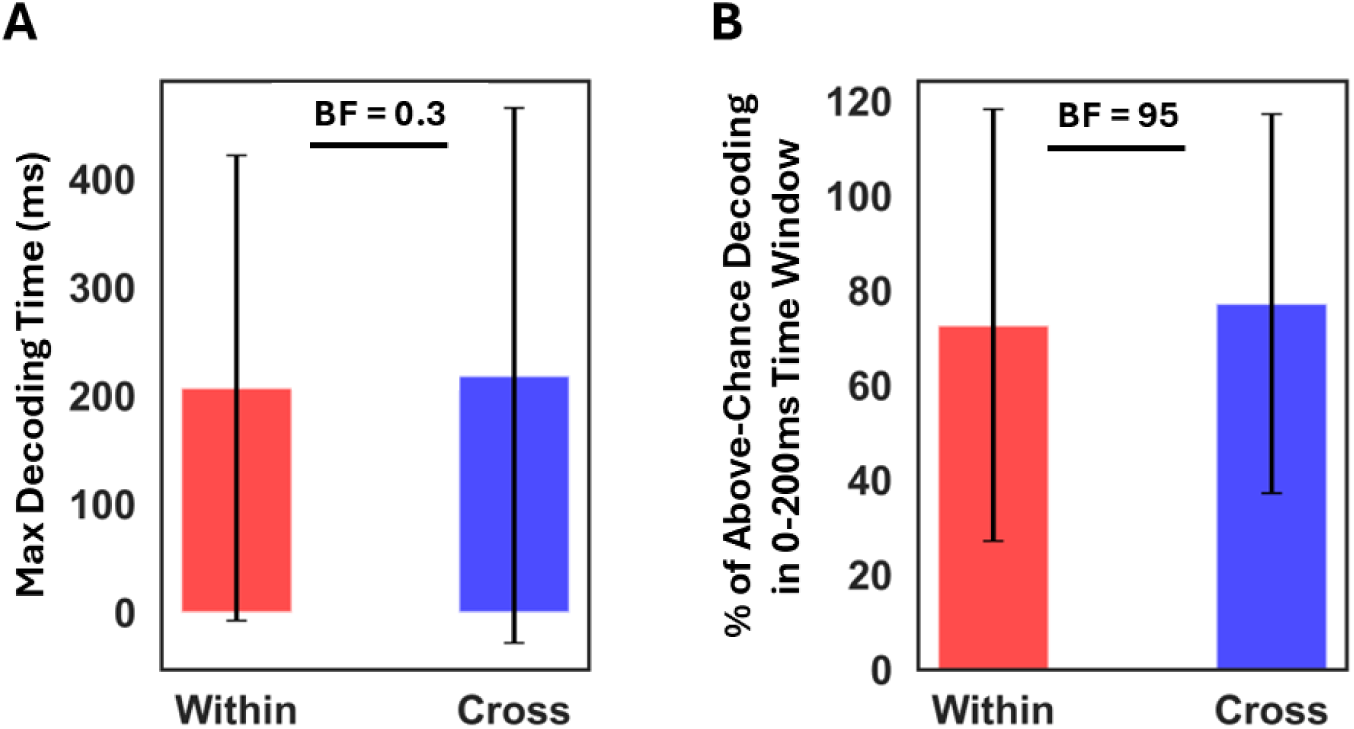
Comparison of the timing of orientation decoding in *within-* and *cross-subject* decoding. **A** shows the time of maximum decoding in the *within-* and *cross-subject* decoding analyses. **B** shows the percentage of above-chance decoding scores found in the earlier time window (0-200ms) in the *within-* and *cross-subject* decoding analyses. Error bars indicate SEM over subjects.

**Supplementary Figure 12.**
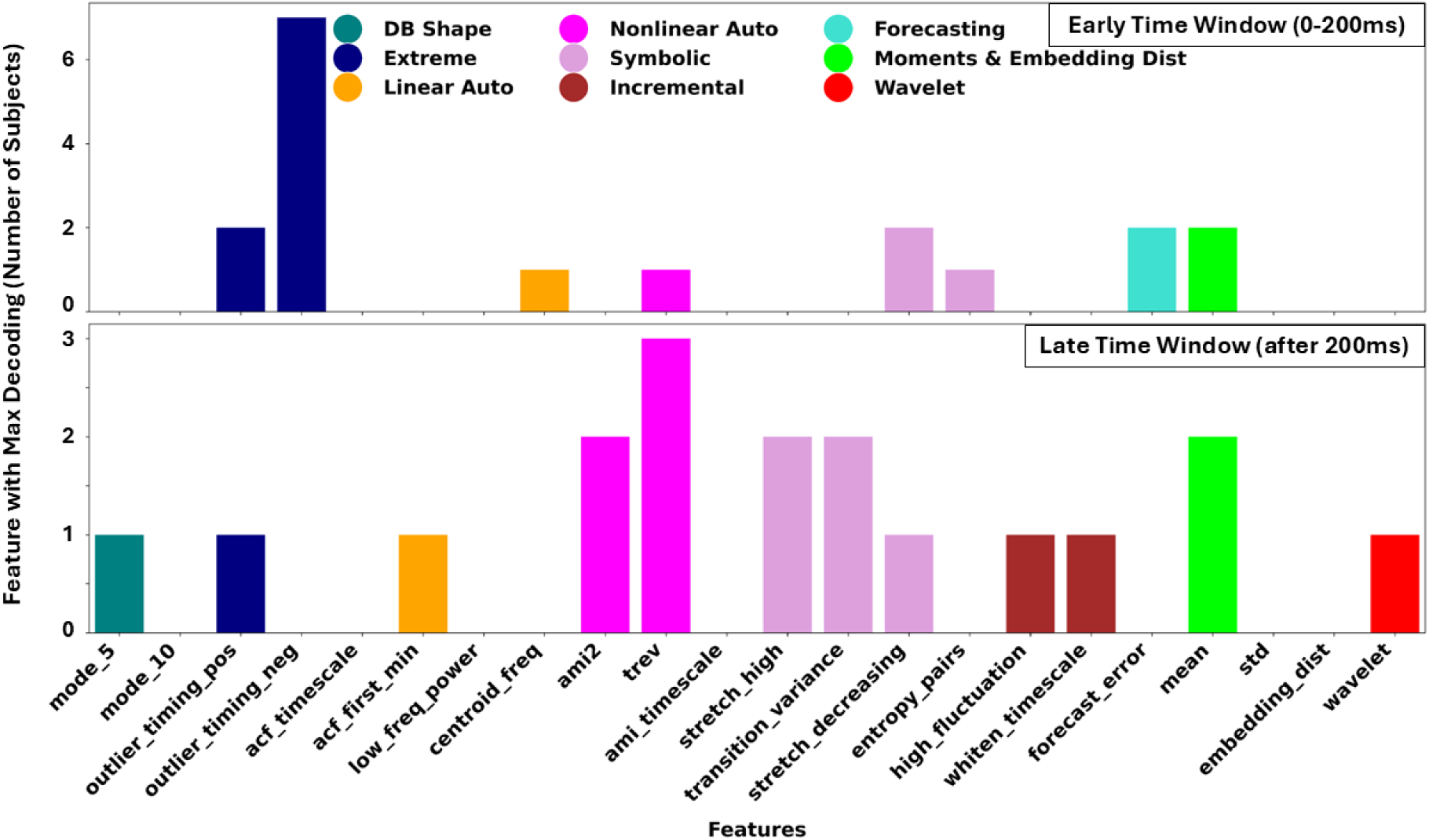
Distribution of features which provided the highest orientation decoding. Top and bottom panels show the results for the earlier (0-200ms) and later (after 200ms) analysis window, respectively.

## Footnotes

1 https://openneuro.org/datasets/ds002550/versions/1.0.^1^

2 https://time-series-features.gitbook.io/Catch22-features/Catch22/the-dn_outlierinclude-features

3 https://github.com/DynamicsAndNeuralSystems/pyCatch22?tab=readme-ov-file

## References

Aggarwal, S., & Chugh, N. (2022). Review of machine learning techniques for EEG based brain computer interface. Archives of Computational Methods in Engineering, 29(5), 3001–3020.

Ahmadian, Y., Pillow, J. W., & Paninski, L. (2011). Efficient Markov chain Monte Carlo methods for decoding neural spike trains. Neural computation, 23(1), 46–96.

Auge, D., Hille, J., Mueller, E., & Knoll, A. (2021). A survey of encoding techniques for signal processing in spiking neural networks. Neural Processing Letters, 53(6), 4693–4710.

Baillet, S. (2017). Magnetoencephalography for brain electrophysiology and imaging. Nature neuroscience, 20(3), 327–339.

Barbieri, R., Frank, L. M., Nguyen, D. P., Quirk, M. C., Solo, V., Wilson, M. A., & Brown, E. N. (2004). Dynamic analyses of information encoding in neural ensembles. Neural computation, 16(2), 277–307.

Barbieri, R., Frank, L., Nguyen, D., Quirk, M., Solo, V., Wilson, M., & Brown, E. (2004). A Bayesian decoding algorithm for analysis of information encoding in neural ensembles. The 26th Annual International Conference of the IEEE Engineering in Medicine and Biology Society.

Barbieri, R., Wilson, M. A., Frank, L. M., & Brown, E. N. (2005). An analysis of hippocampal spatio-temporal representations using a Bayesian algorithm for neural spike train decoding. IEEE transactions on neural systems and rehabilitation engineering, 13(2), 131–136.

Bishop, W., Byron, M. Y., Santhanam, G., Afshar, A., Ryu, S. I., Shenoy, K. V., Vogelstein, R. J., Beaty, J., & Harshbarger, S. (2008). The use of a virtual integration environment for the real-time implementation of neural decode algorithms. 2008 30th Annual International Conference of the IEEE Engineering in Medicine and Biology Society.

Blankertz, B., Lemm, S., Treder, M., Haufe, S., & Müller, K.-R. (2011). Single-trial analysis and classification of ERP components—a tutorial. NeuroImage, 56(2), 814–825.

Borst, A., & Theunissen, F. E. (1999). Information theory and neural coding. Nature neuroscience, 2(11), 947–957.

Breiman, L. (2001). Random forests. Machine learning, 45, 5–32.

Carlson, T. A., Grootswagers, T., & Robinson, A. K. (2019). An introduction to time-resolved decoding analysis for M/EEG. arXiv preprint arXiv:1905.04820.

Chen, W., Wang, Y., Cao, G., Chen, G., & Gu, Q. (2014). A random forest model based classification scheme for neonatal amplitude-integrated EEG. Biomedical engineering online, 13, 1–13.

Cohen, I., Huang, Y., Chen, J., Benesty, J., Benesty, J., Chen, J., Huang, Y., & Cohen, I. (2009). Pearson correlation coefficient. Noise reduction in speech processing, 1-4.

Csaky, R., Van Es, M. W., Jones, O. P., & Woolrich, M. (2023). Group-level brain decoding with deep learning. Human Brain Mapping, 44(17), 6105–6119.

Curto, C., Degeratu, A., & Itskov, V. (2013). Encoding binary neural codes in networks of threshold-linear neurons. Neural computation, 25(11), 2858–2903.

Daubechies, I. (1992). Ten lectures on wavelets. Society for industrial and applied mathematics.

Endres, D. M., Földiák, P., & Priss, U. (2009). An application of formal concept analysis to semantic neural decoding. Annals of Mathematics and Artificial Intelligence, 57, 233–248.

Fazel-Rezai, R., Allison, B. Z., Guger, C., Sellers, E. W., Kleih, S. C., & Kübler, A. (2012). P300 brain computer interface: current challenges and emerging trends. Frontiers in neuroengineering, 5, 14.

Feldman, D. E. (2012). The spike-timing dependence of plasticity. Neuron, 75(4), 556–571.

Fernández-Delgado, M., Cernadas, E., Barro, S., & Amorim, D. (2014). Do we need hundreds of classifiers to solve real world classification problems? The journal of machine learning research, 15(1), 3133–3181.

Fulcher, B. D., & Jones, N. S. (2017). hctsa: A computational framework for automated time-series phenotyping using massive feature extraction. Cell systems, 5(5), 527–531. e523.

Glaser, J. I., Benjamin, A. S., Chowdhury, R. H., Perich, M. G., Miller, L. E., & Kording, K. P. (2020). Machine learning for neural decoding. eneuro, 7(4).

Graf, A. B., Kohn, A., Jazayeri, M., & Movshon, J. A. (2011). Decoding the activity of neuronal populations in macaque primary visual cortex. Nature neuroscience, 14(2), 239–245.

Grootswagers, T., Wardle, S. G., & Carlson, T. A. (2017). Decoding dynamic brain patterns from evoked responses: A tutorial on multivariate pattern analysis applied to time series neuroimaging data. Journal of cognitive neuroscience, 29(4), 677–697.

Gross, J. (2019). Magnetoencephalography in cognitive neuroscience: a primer. Neuron, 104(2), 189–204.

Guger, C., Daban, S., Sellers, E., Holzner, C., Krausz, G., Carabalona, R.,…& Edlinger, G. (2009). How many people are able to control a P300-based brain–computer interface (BCI)?. Neuroscience letters, 462(1), 94–98.

Hämäläinen, M., Hari, R., Ilmoniemi, R. J., Knuutila, J., & Lounasmaa, O. V. (1993). Magnetoencephalography—theory, instrumentation, and applications to noninvasive studies of the working human brain. Reviews of modern Physics, 65(2), 413.

Hancock, J. T., & Khoshgoftaar, T. M. (2020). Survey on categorical data for neural networks. Journal of big data, 7(1), 28.

Hari, R., & Salmelin, R. (2012). Magnetoencephalography: From SQUIDs to neuroscience: Neuroimage 20th anniversary special edition. NeuroImage, 61(2), 386–396.

Harris, K. D. (2005). Neural signatures of cell assembly organization. Nature reviews neuroscience, 6(5), 399–407.

Hatsopoulos, N. G., & Donoghue, J. P. (2009). The science of neural interface systems. Annual review of neuroscience, 32(1), 249–266.

Haxby, J. V., Gobbini, M. I., Furey, M. L., Ishai, A., Schouten, J. L., & Pietrini, P. (2001). Distributed and overlapping representations of faces and objects in ventral temporal cortex. Science, 293(5539), 2425–2430.

Hebart, M. N., Bankson, B. B., Harel, A., Baker, C. I., & Cichy, R. M. (2018). The representational dynamics of task and object processing in humans. Elife, 7, e32816.

Hill, R. M., Schofield, H., Boto, E., Rier, L., Osborne, J., Doyle, C., Worcester, F., Hayward, T., Holmes, N., & Bowtell, R. (2024). Optimising the sensitivity of optically-pumped magnetometer magnetoencephalography to gamma band electrophysiological activity. Imaging Neuroscience, 2, 1–19.

Karimi-Rouzbahani, H. (2024). Evidence for multiscale multiplexed representation of visual features in EEG. Neural computation, 36(3), 412–436.

Karimi-Rouzbahani, H., & McGonigal, A. (2024). Generalisability of epileptiform patterns across time and patients. Scientific Reports, 14(1), 6293.

Karimi-Rouzbahani, H., & McGonigal, A. (2025). Directionality of neural activity in and out of the seizure onset zone in focal epilepsy. Network Neuroscience, 1-26.

Karimi-Rouzbahani, H., Bagheri, N., & Ebrahimpour, R. (2017). Average activity, but not variability, is the dominant factor in the representation of object categories in the brain. Neuroscience, 346, 14–28.

Karimi-Rouzbahani, H., Rich, A. N., & Woolgar, A. (2024). Spatiotemporal characterisation of information coding and exchange in the multiple demand network. bioRxiv, 2024-10.

Karimi-Rouzbahani, H., Shahmohammadi, M., Vahab, E., Setayeshi, S., & Carlson, T. (2021). Temporal variabilities provide additional category-related information in object category decoding: a systematic comparison of informative EEG features. Neural computation, 33(11), 3027–3072.

Karimi-Rouzbahani, H., Vahab, E., Ebrahimpour, R., & Menhaj, M. B. (2019). Spatiotemporal analysis of category and target-related information processing in the brain during object detection. Behavioural brain research, 362, 224–239.

Kartsaki, E., Hilgen, G., Sernagor, E., & Cessac, B. (2024). How Does the Inner Retinal Network Shape the Ganglion Cells Receptive Field? A Computational Study. Neural computation, 36(6), 1041–1083.

Kass, R. E., & Raftery, A. E. (1995). Bayes factors. Journal of the american statistical association, 90(430), 773–795.

Kay, K. N., Naselaris, T., Prenger, R. J., & Gallant, J. L. (2008). Identifying natural images from human brain activity. Nature, 452(7185), 352–355.

Kemere, C., Sahani, M., & Meng, T. (2003). Robust neural decoding of reaching movements for prosthetic systems. Proceedings of the 25th Annual International Conference of the IEEE Engineering in Medicine and Biology Society (IEEE Cat. No. 03CH37439).

Khademi, Z., Ebrahimi, F., & Kordy, H. M. (2023). A review of critical challenges in MI-BCI: From conventional to deep learning methods. Journal of Neuroscience Methods, 383, 109736.

Kietzmann, T. C., Spoerer, C. J., Sörensen, L. K., Cichy, R. M., Hauk, O., & Kriegeskorte, N. (2019). Recurrence is required to capture the representational dynamics of the human visual system. Proceedings of the National Academy of Sciences, 116(43), 21854–21863.

King, J.-R., & Dehaene, S. (2014). Characterizing the dynamics of mental representations: the temporal generalization method. Trends in cognitive sciences, 18(4), 203–210.

Koide-Majima, N., Nishimoto, S., & Majima, K. (2024). Mental image reconstruction from human brain activity: Neural decoding of mental imagery via deep neural network-based Bayesian estimation. Neural Networks, 170, 349–363.

Kutas, M., & Dale, A. (2013). Electrical and magnetic readings of mental functions. In Cognitive neuroscience (pp. 197-242). Psychology press.

Lubba, C. H., Sethi, S. S., Knaute, P., Schultz, S. R., Fulcher, B. D., & Jones, N. S. (2019). Catch22: CAnonical Time-series CHaracteristics: Selected through highly comparative time-series analysis. Data Mining and Knowledge Discovery, 33(6), 1821–1852.

Malik, W. Q., Truccolo, W., Brown, E. N., & Hochberg, L. R. (2010). Efficient decoding with steady-state Kalman filter in neural interface systems. IEEE transactions on neural systems and rehabilitation engineering, 19(1), 25–34.

Miller, M. B., Van Horn, J. D., Wolford, G. L., Handy, T. C., Valsangkar-Smyth, M., Inati, S.,…& Gazzaniga, M. S. (2002). Extensive individual differences in brain activations associated with episodic retrieval are reliable over time. Journal of Cognitive Neuroscience, 14(8), 1200–1214.

Miyawaki, Y., Uchida, H., Yamashita, O., Sato, M.-a., Morito, Y., Tanabe, H. C., Sadato, N., & Kamitani, Y. (2008). Visual image reconstruction from human brain activity using a combination of multiscale local image decoders. Neuron, 60(5), 915–929.

Moerel, M., Yacoub, E., Gulban, O. F., Lage-Castellanos, A., & De Martino, F. (2021). Using high spatial resolution fMRI to understand representation in the auditory network. Progress in neurobiology, 207, 101887.

Mohsenzadeh, Y., Mullin, C., Oliva, A., & Pantazis, D. (2019). The perceptual neural trace of memorable unseen scenes. Scientific reports, 9(1), 6033.

Naselaris, T., Prenger, R. J., Kay, K. N., Oliver, M., & Gallant, J. L. (2009). Bayesian reconstruction of natural images from human brain activity. Neuron, 63(6), 902–915.

Oram, M. W., Földiák, P., Perrett, D. I., & Sengpiel, F. (1998). TheIdeal Homunculus’: decoding neural population signals. Trends in neurosciences, 21(6), 259–265.

Pan, J., Chen, X., Ban, N., He, J., Chen, J., & Huang, H. (2022). Advances in P300 brain–computer interface spellers: toward paradigm design and performance evaluation. Frontiers in human neuroscience, 16, 1077717.

Paninski, L., Pillow, J., & Lewi, J. (2007). Statistical models for neural encoding, decoding, and optimal stimulus design. Progress in brain research, 165, 493–507.

Panzeri, S., Brunel, N., Logothetis, N. K., & Kayser, C. (2010). Sensory neural codes using multiplexed temporal scales. Trends in neurosciences, 33(3), 111–120.

Papadimitriou, C. H., & Friederici, A. D. (2022). Bridging the gap between neurons and cognition through assemblies of neurons. Neural computation, 34(2), 291–306.

Poucet, Β., Lenck-Santini, P., Hok, V., Save, E., Banquet, J., Gaussier, P., & Muller, R. (2004). Spatial navigation and hippocampal place cell firing: the problem of goal encoding. Reviews in the Neurosciences, 15(2), 89–108.

Quentin, R., King, J.-R., Sallard, E., Fishman, N., Thompson, R., Buch, E. R., & Cohen, L. G. (2019). Differential brain mechanisms of selection and maintenance of information during working memory. Journal of Neuroscience, 39(19), 3728–3740.

Rajaei, K., Mohsenzadeh, Y., Ebrahimpour, R., & Khaligh-Razavi, S. M. (2019). Beyond core object recognition: Recurrent processes account for object recognition under occlusion. PLoS computational biology, 15(5), e1007001.

Rajpura, P., Cecotti, H., & Meena, Y. K. (2024). Explainable artificial intelligence approaches for brain-computer interfaces: a review and design space. Journal of Neural Engineering.

Ritchie, J. B., Tovar, D. A., & Carlson, T. A. (2015). Emerging object representations in the visual system predict reaction times for categorization. PLoS computational biology, 11(6), e1004316.

Rybakken, E., Baas, N., & Dunn, B. (2019). Decoding of neural data using cohomological feature extraction. Neural computation, 31(1), 68–93.

Santos-Mayo, A., Gilbert, F., Ahumada, L., Traiser, C., Engle, H., Panitz, C., Ding, M., & Keil, A. (2025). Decoding in the fourth dimension: Classification of temporal patterns and their generalization across locations. Human Brain Mapping, 46(2), e70152.

Schalk, G., McFarland, D. J., Hinterberger, T., Birbaumer, N., & Wolpaw, J. R. (2004). BCI2000: a general-purpose brain-computer interface (BCI) system. IEEE Transactions on biomedical engineering, 51(6), 1034–1043.

Schofield, H., Hill, R. M., Feys, O., Holmes, N., Osborne, J., Doyle, C., Bobela, D., Corvilian, P., Wens, V., & Rier, L. (2024). A Novel, Robust, and Portable Platform for Magnetoencephalography using Optically Pumped Magnetometers. bioRxiv.

Siegel, M., Donner, T. H., & Engel, A. K. (2012). Spectral fingerprints of large-scale neuronal interactions. Nature Reviews Neuroscience, 13(2), 121–134.

Taghizadeh-Sarabi, M., Daliri, M. R., & Niksirat, K. S. (2015). Decoding objects of basic categories from electroencephalographic signals using wavelet transform and support vector machines. Brain topography, 28, 33–46.

Tam, W.-k., Wu, T., Zhao, Q., Keefer, E., & Yang, Z. (2019). Human motor decoding from neural signals: a review. BMC Biomedical Engineering, 1, 1–22.

Thorpe, S., Fize, D., & Marlot, C. (1996). Speed of processing in the human visual system. Nature, 381(6582), 520–522.

van Gerven, M. A., Seeliger, K., Güçlü, U., & Güçlütürk, Y. (2019). Current advances in neural decoding (pp. 379-394). Springer International Publishing.

Vargas-Hakim, G.-A., Mezura-Montes, E., & Acosta-Mesa, H.-G. (2021). A review on convolutional neural network encodings for neuroevolution. IEEE Transactions on Evolutionary Computation, 26(1), 12–27.

Wang, S., Liu, S., Tan, Z., & Wang, X. (2024). Mindbridge: A cross-subject brain decoding framework. Proceedings of the IEEE/CVF Conference on Computer Vision and Pattern Recognition.

Wang, X., Wirth, A., & Wang, L. (2007). Structure-based statistical features and multivariate time series clustering. Seventh IEEE international conference on data mining (ICDM 2007), (pp. 351-360). IEEE.

Wei, C.-S., Keller, C. J., Li, J., Lin, Y.-P., Nakanishi, M., Wagner, J., Wu, W., Zhang, Y., & Jung, T.-P. (2021). Inter-and intra-subject variability in brain imaging and decoding. In (Vol. 15, pp. 791129): Frontiers Media SA.

Wittevrongel, B., Holmes, N., Boto, E., Hill, R., Rea, M., Libert, A., Khachatryan, E., Van Hulle, M. M., Bowtell, R., & Brookes, M. J. (2021). Practical real-time MEG-based neural interfacing with optically pumped magnetometers. BMC biology, 19(1), 158.

Wolpaw, J. R., Birbaumer, N., McFarland, D. J., Pfurtscheller, G., & Vaughan, T. M. (2002). Brain–computer interfaces for communication and control. Clinical neurophysiology, 113(6), 767–791.

Wu, W., Kulkarni, J. E., Hatsopoulos, N. G., & Paninski, L. (2009). Neural decoding of hand motion using a linear state-space model with hidden states. IEEE transactions on neural systems and rehabilitation engineering, 17(4), 370–378.

Zhang, J., Li, C., Liu, G., Min, M., Wang, C., Li, J., Wang, Y., Yan, H., Zuo, Z., & Huang, W. (2022). A CNN-transformer hybrid approach for decoding visual neural activity into text. Computer Methods and Programs in Biomedicine, 214, 106586.

